# Ultrastructural plasma membrane asymmetries in tension and curvature promote yeast cell fusion

**DOI:** 10.1101/2021.03.18.435973

**Authors:** Olivia Muriel, Laetitia Michon, Wanda Kukulski, Sophie G Martin

## Abstract

Cell-cell fusion is central to the process of fertilization for sexual reproduction. This necessitates the remodeling of peri-cellular matrix or cell wall material and the merging of plasma membranes. In walled fission yeast *S. pombe*, the fusion of P and M cells during sexual reproduction relies on the fusion focus, an actin structure that concentrates glucanase-containing secretory vesicles for local cell wall digestion necessary for membrane fusion. Here, we present a correlative light and electron microscopy (CLEM) quantitative study of a large dataset of 3D tomograms of the fusion site, which revealed the ultrastructure of the fusion focus as an actin-containing, vesicle-dense structure excluding other organelles. Unexpectedly, the data revealed asymmetries between the two gametes: M-cells exhibit a taut and convex plasma membrane that progressively protrudes into P-cells, which exhibit a more slack, wavy plasma membrane. These asymmetries are relaxed upon plasma membrane fusion, with observations of ramified pores that may result from multiple initiations or inhomogeneous expansion. We show that P-cells have a higher exo-to endocytosis ratio than M-cells, and that local reduction in exocytosis abrogates membrane waviness and compromises cell fusion significantly more in P-than M-cells. Reciprocally, reduction of turgor pressure specifically in M-cells prevents their protrusions into P-cells and delays cell fusion. Thus, asymmetric membrane conformations, which result from differential turgor pressure and exocytosis/endocytosis ratios between mating types, favor cell-cell fusion.

## Introduction

Cell-cell fusion is a fundamental process, which underlies sexual reproduction and also occurs between somatic cells during the development of various tissues. The successful merging of plasma membranes to fuse two cells into one requires a number of preparatory steps. These include the formation of cell adhesion and the remodeling of extracellular material to allow membrane contact. Significant force is also required to overcome repulsive membrane and hydration charges. In some cases, the actin cytoskeleton generates this force and allows the engagement of membrane fusogenic machineries [1, 2]. Fusogenic proteins, which have only been identified in some organisms and cell types [3, 4], then likely drive the formation of fusion pore(s), though this has not been directly observed in any system.

Though fusion processes and machineries are extraordinarily diverse across the tree of life, the two cells engaging in fusion can be described as asymmetric in the vast majority of known fusion processes. Fertilization events largely happen between gametes of widely differing shape and size. Morphological and functional asymmetries have also been described during the fusion of somatic cells. One of the best-described cases is that of muscle development in *Drosophila*, where fusion-competent myoblasts form a podosome that protrudes into the myotube [5]. Myoblast and myotube assume different roles and morphologies, with actin assembly thought to power podosome protrusion, while a myosin II-dependent response in the myotube increases cortical tension to promote fusion [6, 7]. Similar asymmetric cell fusion structures have been described in other types of somatic cell fusion [4]. By contrast, cell fusion during sexual reproduction in yeast is thought to happen between two isogametes, i.e. gametes that do not exhibit any overt morphological distinction.

The differences between yeast gametes are defined by the genetic information present in the active mating type locus, which principally controls the expression of distinct pheromone and receptor genes. Pheromone-dependent communication underlies sexual differentiation and the formation of cell pairs [8]. Wildtype fission yeast cells are naturally homothallic, i.e. they can switch between mating types, making the population mating-competent when faced with nitrogen starvation. Cells can also be fixed in one or other heterothallic mating type, M (*h*-) and P (*h+*), which can mate and fuse when mixed together. The two pheromones have distinct biochemical qualities and modes of secretion, and also exhibit slightly different functions and evolutionary constraints, suggesting some functional diversification between the isogametic mating types [8–10]. However, very few additional genes are differentially expressed between mating types [11, 12], and the cell fusion process itself is largely considered to happen between two symmetric cells.

Yeast cells are encased in a cell wall that needs to be remodeled for cell morphogenesis. For cell growth, which is required for cell pairing during mating, remodeling happens at the cell projection (also known as shmoo) tip, with turgor pressure providing the critical force [13, 14]. For cell fusion, the cell wall needs to be locally digested to allow membrane contact. This requires the formation of an actin-based structure called the fusion focus, which is assembled by the formin Fus1 and serves to concentrate the myosin V-dependent delivery of secretory vesicles at the cell-cell contact site. These vesicles contain hydrolytic enzymes, which are likely locally secreted leading to cell wall erosion [15]. Turgor pressure is hypothesized to then help bring membranes in contact [16]. Interestingly, we had previously described an asymmetry between mating types in the formation of the fusion focus, which is stabilized earlier in M-than P-cells [15]. While advanced live-cell imaging studies have revealed significant molecular and mechanistic information on the assembly of the fusion focus [15, 17–20], how the two cells shape their plasma membranes for merging and how the initial fusion pore expands are not understood.

Here, we used correlative light microscopy and electron tomography to study cell fusion in *S. pombe* at the ultrastructural level. Our data shows the organization of the fusion focus as a vesicle-rich structure that excludes other organelles and describes all fusion stages, including pairs with membrane contact, small and large fusion pores, indicating inhomogeneous pore expansion. Unexpectedly, we reveal that the two pre-fusion gametes exhibit asymmetries in both membrane tension and curvature, with M-cells protruding into P-cells. We trace the origin of these asymmetries to different ratios of endo- and exocytic activities and different turgor pressure between mating types, respectively, and show that these asymmetries promote efficient cell-cell fusion.

## Results

### Ultrastructure of the yeast fusion site

We used correlative light and electron microscopy (CLEM) to study the ultrastructure of the actin fusion focus in homothallic (*h90*) wildtype (WT) strains expressing Fus1-sfGFP and Myo52-tdTomato. Using light microscopy we selected mating pairs with Fus1 and Myo52 signal at the shmoo tip indicating presence of the fusion focus. We subsequently identified these cell pairs in transmission electron microscopy (TEM) and acquired tilt series, from which we generated a data set of 124 three-dimensional electron tomographic reconstructions (Fig S1). Based on the minimal distance between the two plasma membranes (PM) in a mating pair, we assigned each pair to one of four different stages along the fusion process (Fig 1A-D). We distinguish three pre-fusion stages: two early stages where cells are in contact through their cell wall (CW), with an average minimal PM distance of 142 ± 50 nm for the first stage (far-CW-contact; 13 tomograms; Fig 1A), and an average minimal PM distance of 49 ± 26 nm for the second (close-CW-contact; 50 tomograms; Fig 1B, Movie S1), where CW have merged together; and a third stage (PM-contact; 11 tomograms; Fig 1C), with an average minimal PM distance of 5 ± 4 nm, where some regions of CWs are fully digested and the PMs of the partner cells appear in contact. The fourth stage is post-fusion, where fusion pores can be identified and the cytosols of the two cells are continuous. Amongst post-fusion pairs, the vast majority showed a single large pore (43 tomograms; Fig 1D, Movie S2). A further 7 tomograms, which were difficult to classify, showed local loss of density in the cell wall suggestive of continuity between partner cells and are described in more detail below. The distribution of tomograms according to minimal distance between PMs is shown in Fig 1E. While the relative frequency of tomograms in the close-CW-contact, PM-contact and small pore connections may be taken as time proxy for the progression of the fusion process, the earliest (far-CW-contact) and latest stages (large fusion pore) are underestimated, as we did not systematically acquire all tomograms for these stages, which could be readily identified from an initial TEM image at high magnification without tomogram reconstruction.

**Figure 1.**
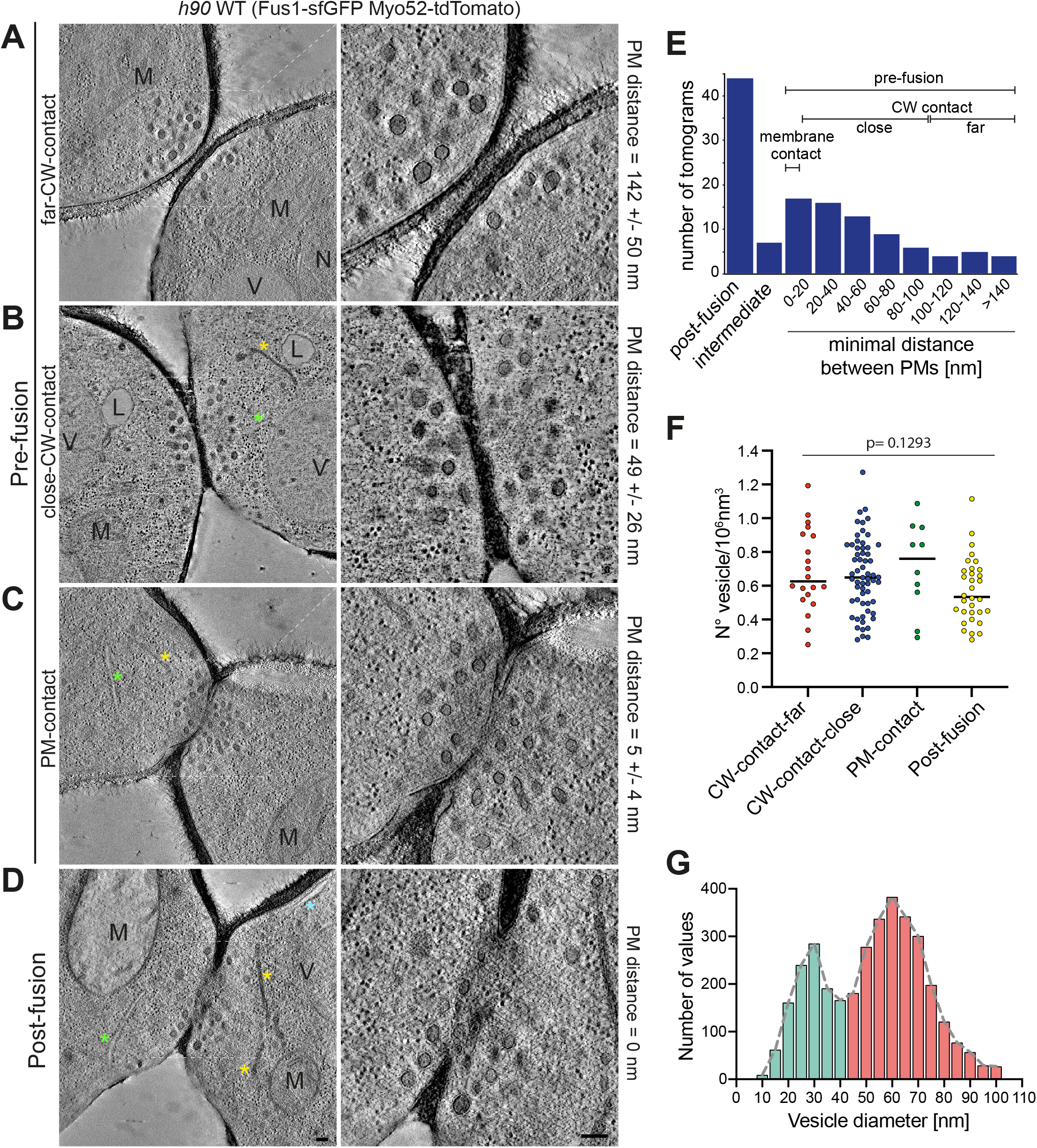
Correlative light-electron analysis of the yeast fusion site. **(A-D)** Virtual z-slices through electron tomograms taken at the contact site of cell pairs during the fusion process as identified by light microscopy (see Fig S1). Examples of different stages in the fusion process are shown: far-CW-contact (A), close-CW-contact (B), PM-contact (C) and post-fusion (D). **(E)** Histogram of the distribution of tomograms according to minimal distance between PMs. **(F)** Density of vesicles within a half-cylinder of diameter 1 μm centred at the centre of the cells’ contact zone (n=20 cells in far-CW-contact; n=60 cells in CW-contact close; n=10 cells in PM-contact; n=32 cells in post-fusion). p=0.1293 (one-way ANOVA test). **(G)** Size distribution of vesicles in cells as in (F) (n=122 cells and 3521 vesicles). Bin width=5 nm. On tomograms, features are marked as following: Blue asterisk = cortical smooth ER; green asterisk = rough ER; yellow asterisk = dense sheet or reticulated organelle; M = mitochondrion; V = vacuole; L = lipid droplet; N = nucleus). Scale bars 100 nm.

Throughout all stages, fusion foci appeared as dense accumulations of dense vesicles. This is expected from the localization of the secretory vesicle-associated type V myosin Myo52 in light microscopy images and from the known co-localization of secretory vesicle markers [15, 17]. We analysed the density and size of the vesicles from 61 randomly chosen tomograms in our dataset (10 far-CW-contact, 30 close-CW-contact, 5 PM-contact, 12 post-fusion and 4 with local loss of density in the cell wall that were included in the postfusion category for this quantification) in a cylinder of defined diameter centred at the cells’ contact (see methods). The density of vesicles did not change along stages (Fig 1F) with an average of 0.65 ± 0.22 vesicles/10^6^ nm^3^. Vesicle size measurements revealed two vesicle populations: the dense vesicles mentioned above with an average diameter of about 60 nm, and a less abundant population of vesicles with an average diameter of about 30 nm. We hypothesize that the larger vesicles are secretory (also confirmed below), whereas the smaller ones represent in part endocytic vesicles, as dimensions are similar to previous measurements in *S. cerevisiae* [21] (Fig 1G). Both vesicle populations were present throughout the different stages, but we noted a somewhat lower relative frequency of the large vesicles in the early stages (Fig S2A). Fusion foci also showed linear structures, likely representing actin filaments, predicted from the activity of the formin Fus1, which co-localized with Myo52 in light microscopy (Fig 2, Movie S1 and S2). While only 51% of tomograms revealed linear filament, which likely reflects the difficulty in preserving and detecting actin structures with this EM technique, the best-quality ones showed a large number of filaments positioned between vesicles and close to the PM. Actin filaments could be seen in both pre- and post-fusion cell pairs and could reach lengths over 0.5μm. Fusion foci were largely devoid of ribosomes or any other organelles, though these were present in other parts of the tomograms (Fig 1A-D, Fig 2). Microtubules were not frequently observed (12% pre-fusion cells), though their frequency increased post fusion (27% post-fusion cells) with microtubules or microtubule bundles crossing the fusion pore, likely in preparation for karyogamy (Fig S2B). Interestingly, 52% cells showed organelles with similar dense appearance to secretory vesicles, which were organized in sheets or reticulated structures (Fig S2C). Though we have not confirmed their identity, this is suggestive of Golgi or endosomal compartments, which may serve for local vesicle production. We note that accumulation of Golgi stacks were also noted in late polarization stages in *S. cerevisiae* mating projections [22]. We conclude that fusion foci are dense vesicle- and actin-rich compartments that cannot easily be penetrated by other large cellular components.

**Figure 2.**
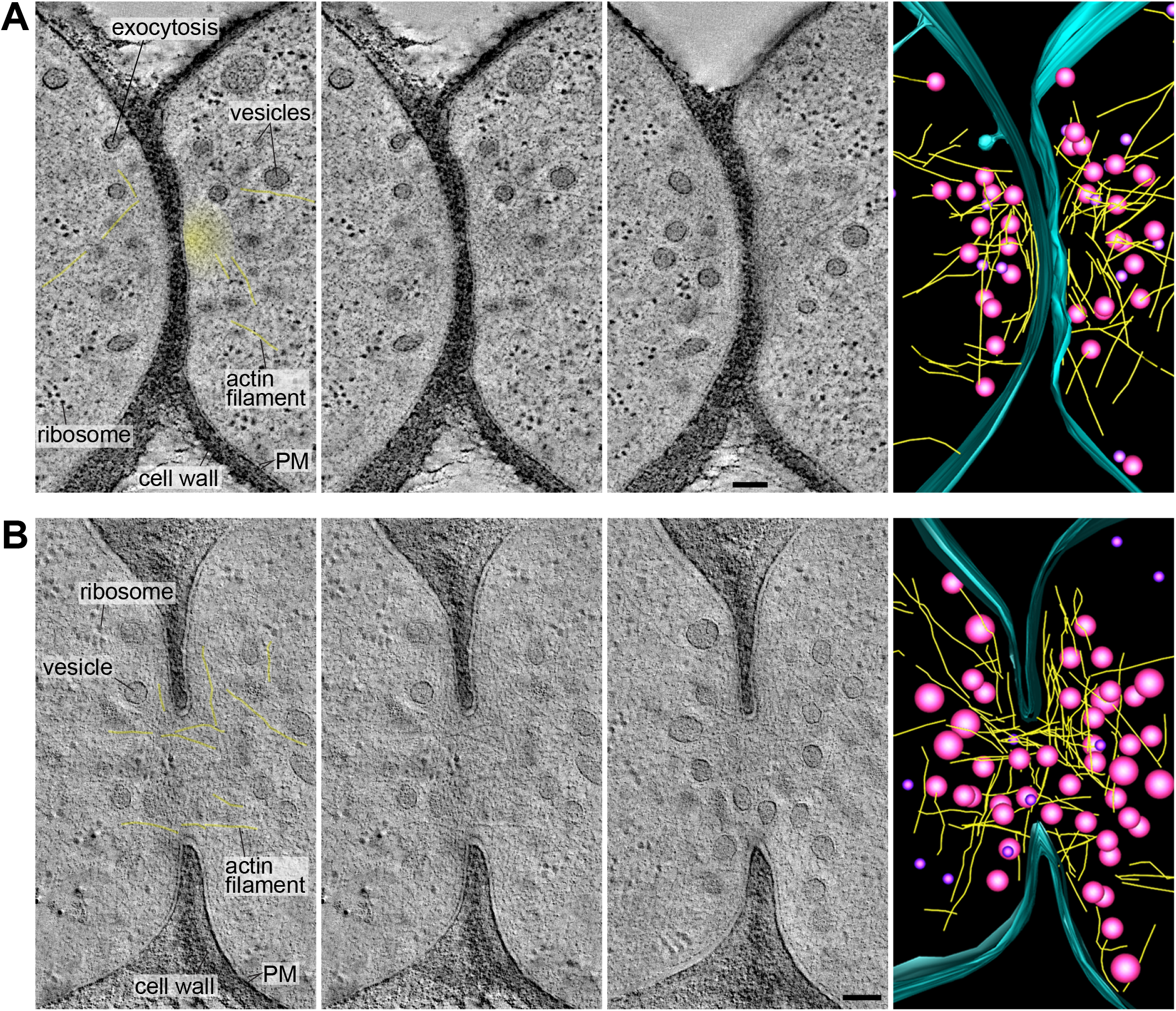
Actin fusion focus ultrastructure. **(A-B)** Virtual z-slices through electron tomograms and model of the actin fusion focus ultrastructure including vesicles (magenta), actin filaments (yellow) and the outer leaflet of the plasma membrane (blue) in cell pairs before (A) and after fusion (B). The first slice is identical to the second and shows labels for the main features in and around the fusion focus with the clearest actin filaments lightly coloured in yellow. The yellow haze in the top pair indicates a region devoid of clear structures, likely dense with actin filaments that cannot be individually resolved. Scale bars 100 nm.

### Shape of the fusion pore(s)

To better understand the membrane merging process, we examined the formation and shape of the fusion pores in the post-fusion cell pairs. We identified a single cell pair with an apparent small pore (of diameter ~45nm) entirely enclosed in the section (Fig 3A). A likely filament can be seen traversing the pore. Seven tomograms showed a local loss of density in the cell wall at the interface between the two cells, suggestive of continuity between the two cells’ cytosols (Fig 3B). However, the inner and outer membrane leaflet could not be unambiguously resolved from the CW density to conclusively distinguish between local cell wall digestion and plasma membrane contact, or possible fusion intermediates or even a fully-formed tiny pore. In three cases, these connections between partner cells were entirely included in the tomographic volume. In the four other cases, they extended to one or both section edges, with one tomogram displaying several such connections, suggesting that membrane merging can initiate at multiple positions (Fig 3C, S3A, Movie S3). However, as these connections extend to the edge of the tomographic volume, we cannot know with certainty whether they initiated independently or represent extensions of a single opening.

**Figure 3.**
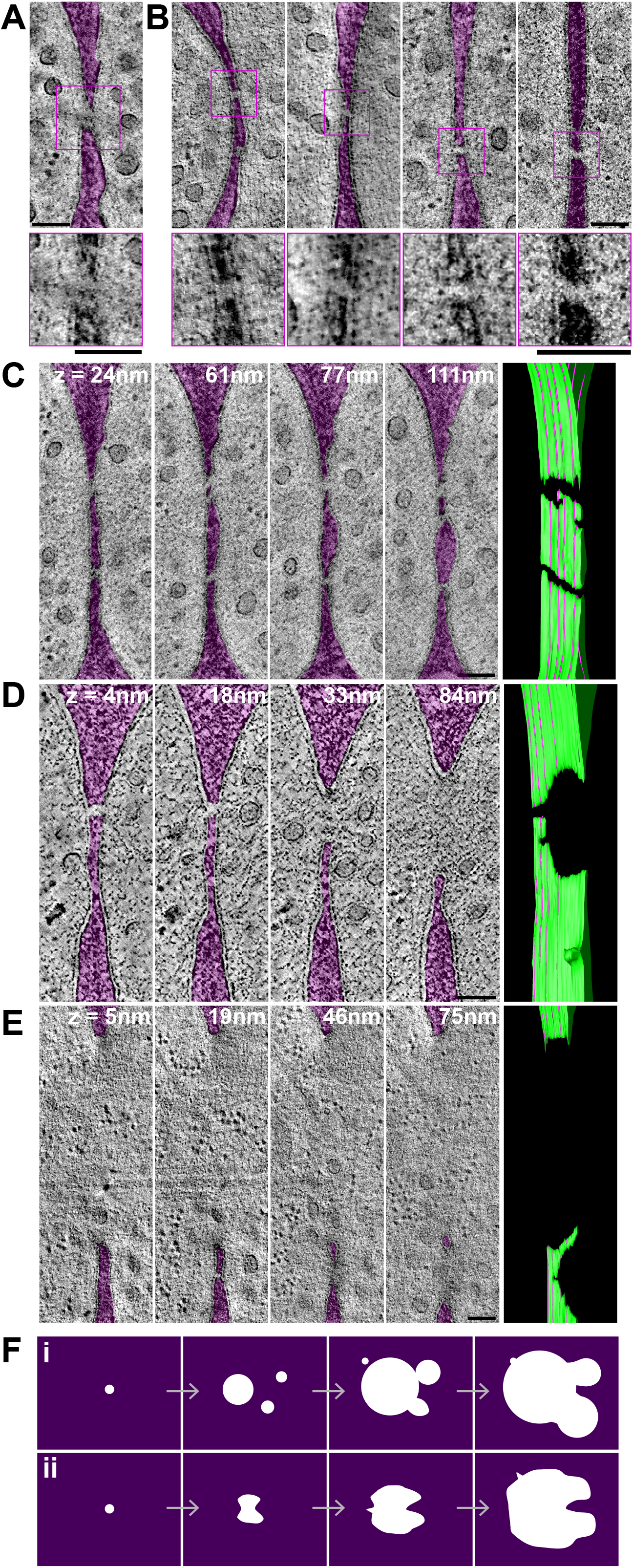
Shape of the fusion pore. **(A-B)** Virtual z-slices through electron tomograms at the contact side of cell pairs, showing one small pore entirely enclosed in the tomographic volume (A) and four cases of apparent connection between cells, with local loss of density in the cell wall, some of which may represent fusion intermediates (B). The cell wall is coloured in transparent purple to help visualization. Zoomed views of the connections without cell wall coloring are shown at the bottom. **(C-E)** Four virtual z-slices through electron tomograms and corresponding model of the outer leaflet of the PM. The location of each z-slice relative to the bottom (left) edge of the model is shown by the purple lines. (C) shows several connections, similar to the ones shown in panel (B). (D) and (E) show large pores with an irregular edge. In (D), the larger pore at z = 84 nm is prolonged into a connection as in (B) at z = 18 and 4 nm. In (E), a large pore at z = 5 nm splits into two openings separated by a strand of cell wall at higher z levels. The cell wall is coloured in transparent purple to help visualization. Images without coloration are shown in Fig S3A-C. **(F)** Schematic drawings for two non-mutually exclusive interpretations about fusion pore formation: initial fusion can occur at several locations forming several small pores that fuse together to form a non-spherical larger pore (i). Fusion initiates at a single location, but pore expansion is limited by cell wall degradation and occurs non-homogeneously, leading to finger-like extensions of the pore periphery (ii). Scale bars 100 nm.

Amongst pairs with a larger fusion pore, two classes retained our attention, as they exhibited non-spherical pore edges. Eight tomograms showed pore edges that extended into irregular extensions, such that at some cross-section of the tomographic volume they appeared as the fusion-intermediates described above (Fig 3D, S3B, S3D, Movie S4-5). Six further tomograms showed a large pore that splits into two openings separated by a strand of cell wall (Fig 3E, S3C, Movie S6). These observations suggest two non-mutually exclusive interpretations. The first is that initial fusion can occur at several locations forming several small pores that fuse together to form a non-spherical larger pore (Fig 3F, i). The second is that fusion usually initiates at a single location, but pore expansion is limited by cell wall degradation and occurs non-homogeneously, leading to finger-like extensions of the pore periphery (Fig 3F, ii). In either case, the small number of tomograms showing membrane contact and small pores suggests that membrane fusion and initial pore expansion are rapid relative to cell wall digestion.

### Plasma membrane shape changes during the fusion process

To learn more about the possible biophysical changes occurring for PM merging, we focused our attention on the conformation of the PM in pre-fusion cell pairs. While yeast gametes are considered isogametes, we identified two morphological asymmetries at the ultrastructural level. First, the membrane at the shmoo tip appeared to exhibit different levels of tension: its appearance ranged from taut, with the membrane forming a smooth arc along the cell wall, to slack/floppy, with the membrane forming waves. Through visual inspection we assigned cells to either the ‘smooth PM’ (sPM) or the ‘wavy PM’ (wPM) category. In the majority of pre-fusion pairs, one cell had wPM and the other sPM (67.5%; 50/74), but other configurations (17.5% or 13/74 with both cells sPM; 15% or 11/74 with both wPM) were also observed (Fig 4A-C). Overall, the frequency of PM waviness, whether in one or both partner cells, increased along the fusion process, with a maximum at the PM-contact stage, and decreased post-fusion (Fig 4D). Indeed, most post-fusion pairs showed sPM (72%; 31/43), with only few pairs showing wPM in one (12%; 5/43) or both (14%; 6/43) cell sides (1 of the 43 could not be clearly classified).

**Figure 4.**
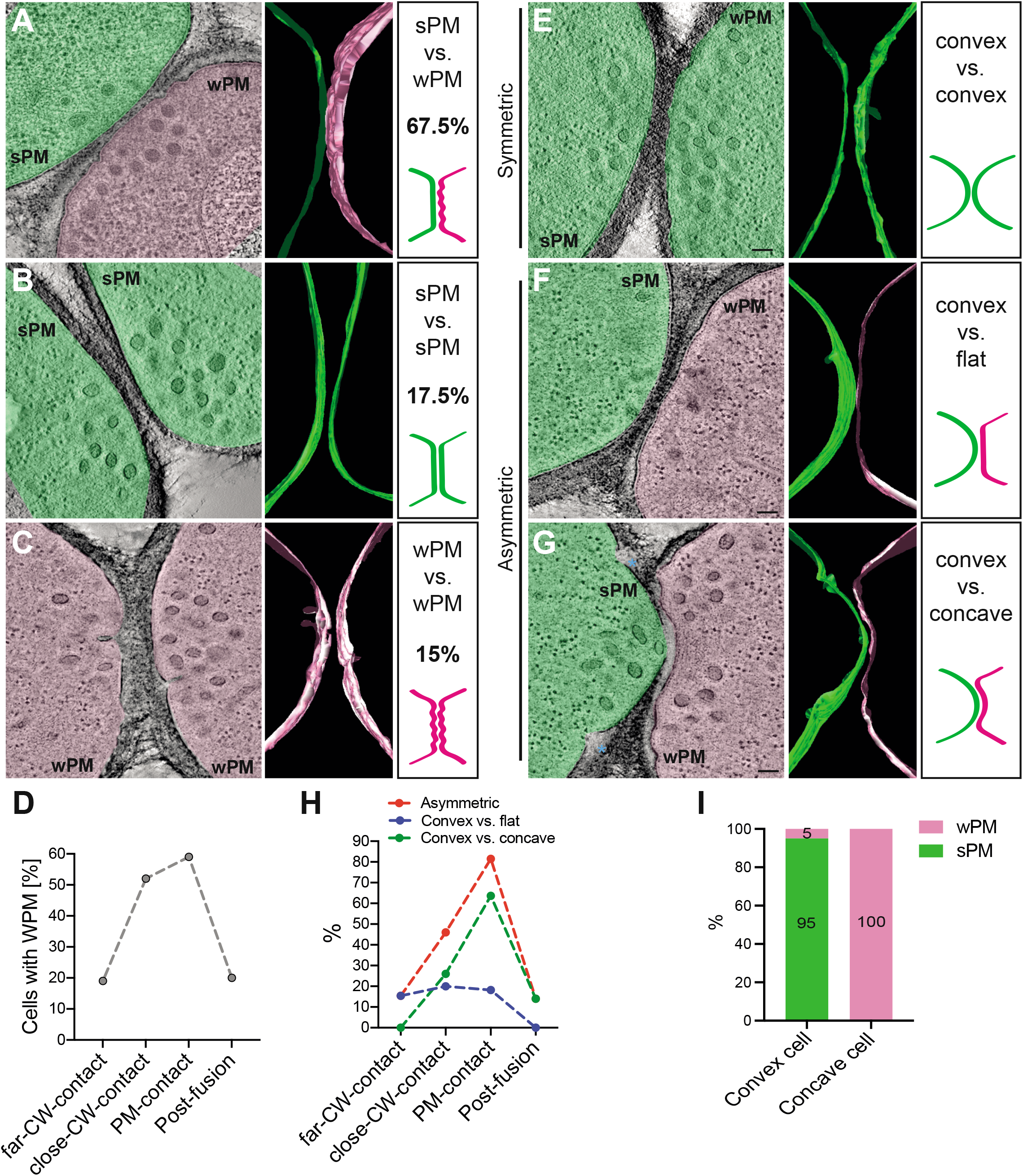
Plasma membrane shape changes in preparation for fusion. **(A-C)** Virtual z-slices through representative electron tomograms of mating cells showing smooth plasma membrane (sPM) versus wavy plasma membrane (wPM) (the predominant situation; A), sPM in both (B) and wPM in both cells (C). Segmentation models of the outer leaflet of the PM and schematic drawings are shown at the right of each example. **(D)** Percentage of cells with wPM along different stages of fusion (n=26 cells for far-CW-contact; n=100 for close-CW-contact; n=22 for PM-contact and n=86 for post-fusion). **(E-G)** Virtual slices through representative electron tomograms of mating cells showing symmetric (convex vs. convex; E) and asymmetric PM curvatures (convex vs. flat (F) and convex vs. concave (G)). Segmentation models of the outer leaflet of the PM and schematic drawing are shown at the right of each example. **(H)** Percentage of asymmetric mating pairs along the different stages of mating (n=13 pairs far-CW-contact; n=50 pairs close-CW-contact; n=11 pairs PM-contact and n=43 pairs post-fusion). **(I)** Percentage of cells with sPM or wPM in pre-fusion pairs showing a convex-concave configuration, showing that low PM tension is associated with the concave state. n=20 pairs. Scale bars 100 nm.

Second, we found that the curvature of the zone of cell-cell contact varied, ranging from convex to flat to even concave. When both partner cells showed similar positive or null curvature (both convex or both flat), we called this situation symmetric (Fig 4E). When the two cells showed distinct curvature, we called this situation asymmetric (Fig 4F-G). Asymmetry was observed in 46% (34/74) of pre-fusion cell pairs. In these pairs, one cell exhibits positive curvature (convex) while the other is either flat (Fig 4F) or exhibits negative curvature (concave) (Fig 4G), such that one cell protrudes into the other one (see also Fig S4). Several protruding cells also showed unresolved membrane invaginations on either side of the protrusion, of unknown origin (Fig S4). The frequency of asymmetric pairs, especially convex vs. concave, increased along the fusion process, with a maximum >80% at the PM-contact stage, and decreased post-fusion (Fig 4H).

Finally, we found that membrane waviness and overall curvature were highly correlated in pre-fusion cells: 100% of concave cells were wavy and over 90% of convex cells were smooth (Fig 4F-G, 4I, S4). Together, these data show that partner cells become highly asymmetric during the fusion process: one cell exhibits a taut, smooth PM and protrudes into its partner, which exhibits a slack, wavy PM. This asymmetry is relaxed post-fusion.

### Plasma membrane asymmetries are mating type specific

The asymmetric appearance of pre-fusion partner cells suggests that each morphology may be associated with a specific mating type. To address this hypothesis, we repeated our CLEM-tomography approach using WT heterothallic strains (*h+* and *h*-), of which only one was labelled with mCherry-D4H, a PM-localized sterol biosensor [23], to identify mating type, and with Fus1-sfGFP to identify pairs in fusion. We generated a set of 23 tomograms (15 with *h+* labelled, 8 with *h*- labelled) at the CW-contact stage. As in the *h90* strain, we observed PM asymmetries in waviness and curvature: 65% of pairs (15/23) showed sPM vs. wPM and 61% (14/23) showed asymmetric curvature (Fig S5A). Importantly, correlation of tomograms with the light microscopy images showed a strong correlation between cell morphology and mating type: 78% (18/23) of *h+* cells, but only 30% (7/23) of *h*- cells, had wPM (Fig 5A, S5B; the sum is >100% due to cases in which both partner cells have wPM); furthermore, in pairs with asymmetric curvature, 79% (11/14) of the convex cells were *h*- (Fig 5B, S5C). We conclude that cell asymmetries are largely conditioned by mating type, in which the *h*- cell protrudes into an *h+* cell with wavy PM.

**Figure 5.**
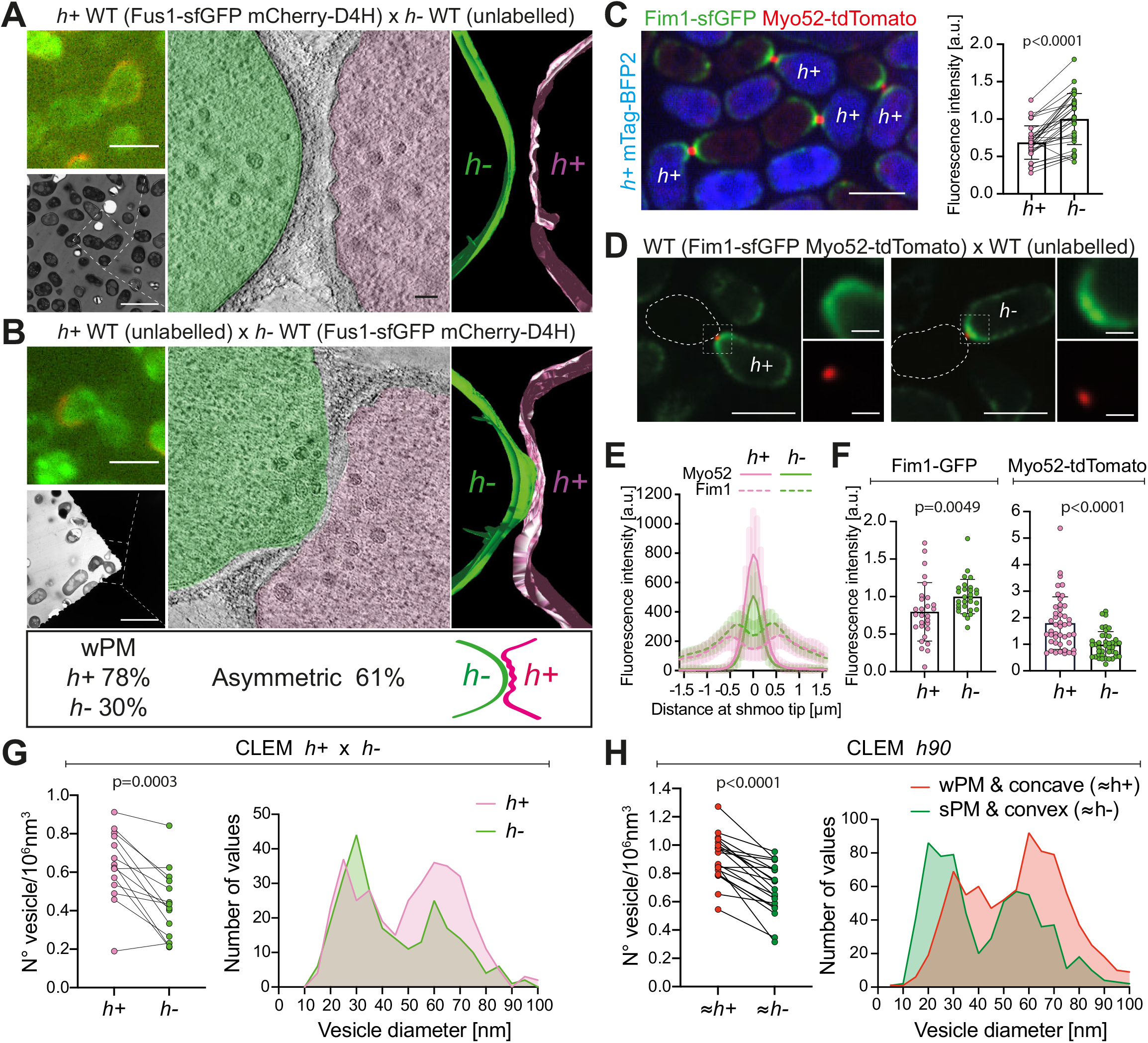
Mating type-specific plasma membrane asymmetries and vesicle distributions. **(A-B)** Representative examples of wildtype cell pairs with one mating type labelled showing wPM versus sPM (A), and convex versus concave PM curvature (B). Top-left light microscopy images show Fus1 in green and cell identity in red, based on the expression of mCherry-D4H. Bottom-left low magnification TEM pictures show the correlation with the same mating pairs. A virtual z-slice of the electron tomogram shows PM shapes, segmented with Imod. Scale bars are 5 μm in light microscopy, 10 μm in low magnification TEM pictures and 100 nm in tomo-grams’ slices. **(C)** Light microscopy images resulting from the average projection of time-lapse movies acquired a maximum speed during 3 minutes. Fim1-sfGFP (green) and Myo52-td-Tomato (red) are expressed in both mating cells. Cytosolic mTag-BFP2 (blue) is expressed only in h+ cells. Paired Fim1 fluorescence intensities at the shmoo tips are shown on the right (n=31 mating pairs in 2 experiments; p<0.0001; paired t-test). Individual data, mean and stdev are shown. Scale bar is 5 μm. **(D)** Light microscopy images resulting from the average projection of time-lapse movies acquired a maximum speed during 3 minutes. Fim1-sfGFP (green) and Myo52-tdTomato (red) are both present only in one cell type (h+ in left images and h-in right images) mated with unlabelled partner. Scale bar are 5 μm for full views and 1 μm for insets. **(E)** Fluorescence profile of Fim1 and Myo52 along the shmoo tips of cells as in (D) (n=13 h+ cells and 13 h-cells). **(F)** Fim1 and Myo52 mean fluorescence intensity of the 5 maximum pixels around each peak from profiles as in (E) (n=28 h+ cells and 29 h-cells from 2 experiments for Fim1 and n=58 h+ cells and 52 h-cells from 4 experiments for Myo52; p values from Mann-Whitney tests). **(G)** Paired density and size distribution of vesicles at the shmoo tips of pre-fusion WT h+/h-cell pairs. n=14 tomograms (264 vesicles for h- and 350 for h+), p=0.0003; paired t-test. **(H)** Paired density and size distribution of vesicles at the shmoo tips of pre-fusion WT h90 cells classified according to their PM curvature and waviness. n=20 tomograms (669 vesicles for sPM/convex and 830 for wPM/concave cells), p<0.0001; paired t-test.

### Cell type-specific balance of exo- and endocytosis

The taut vs floppy PM appearance in *h*- and *h+* cells suggests a possible change in the balance of exo- and endocytosis in the two cell types. Specifically, we hypothesized that PM waves may result from reduced endocytic and/or increased exocytic rates, leading to a steady-state ‘excess’ of PM. To test this hypothesis, we first used fluorescence microscopy to measure the relative levels of proteins associated with endo- and exocytic vesicles. The zone decorated by the endocytic actin patch-associated Fimbrin (Fim1) in pairs with a Myo52-tdTomato-labelled fusion focus was sufficiently internal to unambiguously distinguish the signal from each partner cell. This allowed to compare Fim1-sfGFP levels in *h+* and *h*- partners of the same mating pair, therefore at the same fusion stage, thus eliminating potential stage-specific variations. To distinguish mating types, *h+* cells further expressed cytosolic mTag-BFP2. To average the signal of discrete, dynamic endocytic patches, we measured the average intensity of Fim1-sfGFP signal acquired over 3 minutes. This revealed a 33 ± 12 % reduction in Fim1-sfGFP levels in *h+* cells (Fig 5C), indicating that the amount of endocytosis is lower in *h+* than *h*- cells.

To estimate the amount of secretion we used Myo52-tdTomato, which associates with secretory vesicles. In this case, the fluorescence signals from the two partner cells overlap, preventing direct comparison in a single mating pair. We therefore analysed the signal of Myo52-tdTomato expressed in a single mating type crossed with unlabelled cells of the other mating type. We observed a roughly 2-fold increase in Myo52-tdTomato fluorescence in *h+* cells (Fig 5D-F). As a control, these cells also expressed Fim1-sfGFP, which reproduced the reduced endocytic signal in *h+* cells observed in the direct partner cell comparison, thus validating our results and excluding artifacts due to stage-specific or experimental signal variation. Similar results were obtained when comparing the signal intensity of exocyst subunit Exo84, which associates with secretory vesicles with a 1.5-fold signal increase in *h+* cells (Fig S5D). Thus, *h+* cells show higher levels of exocytic and lower levels of endocytic signal than *h*- cells.

We reasoned that such a difference in vesicular trafficking between mating types should also be observed in the amount and size of vesicles detected by electron microscopy. Indeed, in the labelled *h+* x *h*- cell pairs, the density of vesicles was about 30% higher in *h+* than in *h*- (Fig 5G, left). Furthermore, vesicle size distribution showed that the increased vesicle population in *h+* corresponds to an increased number of 60 nm, secretory vesicles, while the 30 nm vesicle population is slightly reduced (Fig 5G, right). In our larger *h90* tomogram dataset, cell type identity is unknown. However, we reasoned that cell type identity can be inferred in the pairs showing clear PM asymmetries, i-e a convex cell with sPM (inferred to be *h*-) mating with a concave partner with wPM (inferred to be *h+*). In this analysis, we obtained an approximately 23% reduction in the density of vesicles in the convex/sPM cells relative to the concave/wPM (Fig 5H, left). Moreover, the concave/wPM (likely *h+*) cells showed a larger population of 60 nm, secretory vesicles and slight reduction in the 30 nm population (Fig 5H, right). These observations are in line with the fluorescence measurements. Together, this set of data suggests that the difference in plasma membrane waviness between *h+* and *h*- cells is a consequence of altered balance between endocytosis and exocytosis in the two cell types, likely resulting in differences in membrane tension.

### Plasma membrane waviness requires local exocytic activity

To test the idea that membrane waviness is a consequence of excess exocytosis over endocytosis, we aimed to reduce local exocytosis. Deletion of Fus1, necessary for the assembly of the actin fusion focus, is predicted to prevent the strong accumulation of secretory vesicles at the shmoo tip, and thus reduce secretion locally. Because cell fusion completely fails when both partner cells lack *fus1*, we performed our CLEM approach in heterothallic crosses between unlabelled *fus1△* and WT cells expressing Fus1-sfGFP and mCherry-D4H, allowing identification of pairs in fusion and mating type, respectively. Because membrane waviness is mainly observed in *h+* cells, we concentrated most of this analysis on *h+ fus1Δ* x *h-WT* crosses (Fig 6A; n = 14 tomograms), but also obtained some tomograms for *h-fus1Δ* x *h+ WT* crosses (Fig 6B; n = 9 tomograms). As expected, the density of vesicles was strongly reduced in the *fus1△* cells, whether this was *h+* or *h*- (Fig 6A-C). As in previous analysis, the WT *h+* cell showed slightly higher vesicle density than the *h*- cell, which was reduced to similar level in *fus1Δ*. Vesicle size distribution in the WT x *fus1Δ* cross further showed that the reduced vesicle density in *fus1Δ* is due to a strong decrease of the 60 nm vesicles, confirming that these are secretory (Fig 6D). We also searched for linear structures, which we presumed above to be actin filaments: in the 6 tomograms in which we could identify filaments in the WT cell, only one *fus1Δ* cell showed filaments near the cell-cell contact region. These results confirm at the ultrastructural level that Fus1 is necessary for formation of the actin fusion focus and the concentration of secretory vesicles. Examination of secretory vesicle markers Myo52 and Exo84 by light microscopy also showed strong signal reduction (Fig 6E, S5D). Curiously, the signal intensity of the endocytic vesicle marker Fim1-sfGFP was also slightly reduced in *fus1Δ* (Fig 6F). Neither Myo52 nor Fim1 levels were significantly different between mating types (Fig 6E-F). We conclude that local secretion at the cell-cell contact site is decreased in *fus1Δ* cells, and that this likely has consequences on the amount of endocytic activity.

**Figure 6.**
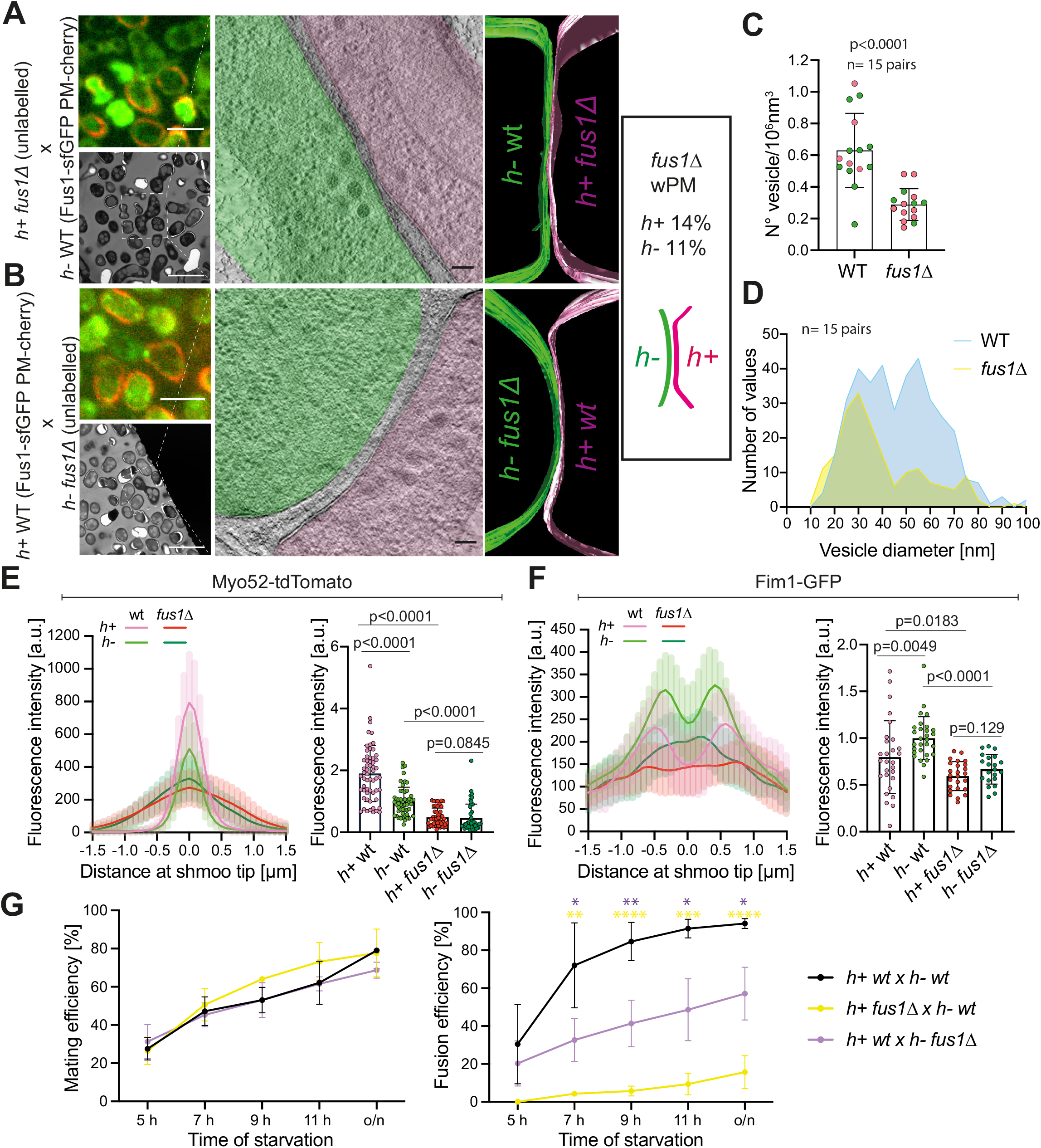
Local exocytosis underlies PM waviness. **(A-B)** Representative example of WT cells crossed with *fus1Δ* cells. The top-left light microscopy pictures show Fus1-sfGFP (green) and mCherry-D4H (red) expressed in the WT cell. Bottom-left low magnification TEM picture show correlation with the same mating pair. The virtual tomogram slices show PM shapes, segmented with Imod. Most *fus1△* cells, whether h+ (A) or h-(B) exhibit sPM. Scale bars are 5 μm in light microscopy, 10 μm in low magnification TEM pictures and 100 nm in tomograms’ slices. **(C)** Density of vesicles at the shmoo tips of WT and *fus1△* pre-fusion cells. n=30 cells, p<0.0001 unpaired t-test. Magenta dots are h+ cells; green dots are h-cells. **(D)** Size distribution of vesicles from (C) (n=30 cells and 379 vesicles for WT and 186 for fus1Δ). Bin width=5. **(E)** Fluorescence profiles of Myo52-tdTomato along the shmoo tips of h+ and h-*fus1△* cells crossed to unlabelled WT on the average projections from 3 min movies at maximum speed (n=13 h+ WT, 13 h-WT, 12 h+ fus1Δ and 9 h-fus1Δ). Right: Myo52 mean fluorescence intensity of the 5 maximum pixels from profiles as on the left (n=51 h+ *fus1Δ* cells and 45 h-*fus1Δ* cells from 4 experiments; p-values are from Mann-Whitney tests). Traces and values for WT, shown for comparison, are identical to those shown in Fig 5E-F. **(F)** Fluorescence profiles of Fim1-sfGFP along the shmoo tips of h+ and h-*fus1Δ* cells on the average projections from 3 min movies at maximum speed (n=13 h+ WT, 13 h-WT, 12 h+ *fus1Δ* and 9 h-*fus1Δ*). Right Fim1 mean fluorescence intensity of 5 maximum pixels from each peak of profiles as on the left (n=28 h+ WT cells, 29 h-WT cells, 25 h+ *fus1Δ* cells and 21 h-*fus1Δ* cells from 2 experiments; p-values from Mann-Whitney tests). Traces and values for WT, shown for comparison, are identical to those shown in Fig 5E-F. **(G)** Quantification of mating and fusion efficiencies over time (n ≥ 500 cells of each genotype from 3-5 experiments; bars show stdev; * indicates p < 0.05; ** indicate p < 0.01; *** indicate p < 0.005; **** indicate p < 0.0001 from unpaired t-tests).

Interestingly, the reduction in local secretion correlates with a strong loss of the wPM phenotype (Fig 6A-B, S6A-B): only 14% of *h+fus1△* cells (2/14) showed wPM, whereas 78% (18/23) and 55% (5/9) of *h+ WT* showed wPM in WT x WT and WT x *fus1△* crosses, respectively. The frequency was similarly reduced in *h*- cells (1/9 vs 7/23). These results are in line with the view that a higher exo-to endocytosis ratio, particularly prominent in *h+* cells, is the cause of the PM waviness.

Regarding PM curvature, our data may suggest some effect of *fus1* deletion. The contact region between cells was wider than in WT crosses with PMs of partner cells parallel over large surfaces (Fig 6A-B, S6A-C), consistent with a fusion delay. The convex cell of asymmetric pairs was usually the WT one (4/5 when WT is *h*-; 3/5 when WT is *h+*) and WT cells were clearly able to protrude into *fus1△* cells (2 *h*- and 1 *h+* examples). However, we also recovered one pair where the *h+fus1Δ* cell appeared to protrude into *h*- WT, though this sample was unfortunately poorly preserved during EM preparation. Our dataset is unfortunately too small to make firm conclusions on the role of Fus1 for cell protrusion.

Given the prominence of the wPM phenotype in the *h+* cell type, it is interesting to note that previous measurements of fusion efficiencies in *fus1Δ* x WT crosses suggested that the defect is more pronounced when *fus1* is deleted in the *h+* cell [15]. We indeed confirmed this observation: while the ability to form cell pairs was not impaired upon deletion of *fus1* in one or the other cell type, fusion efficiency was significantly more strongly compromised when the *h+* cell was *fus1Δ* (Fig 6G). While the main consequence of the loss of focalized secretion in *fus1Δ* may be the inability to digest the cell wall, we hypothesize that loss of the wPM contributes to the enhanced fusion failure in *h*- WT x *h+ fus1Δ* cell pairs.

### Cell protrusion requires strong turgor pressure

A second hypothesis for the origin of PM asymmetries is that they may result from differences in turgor pressure in the two cell types. Specifically, higher turgor in the *h*- cell may enhance membrane tension and facilitate protrusion into the *h+* cell.

To start testing this hypothesis we made use of the *gpd1Δ* mutant. Gpd1 is a glycerol-3-phosphate dehydrogenase necessary for glycerol production to restore turgor pressure in response to osmotic stress [24, 25]. Cells lacking *gpd1* exhibit reduced growth force when challenged in hyper-osmotic conditions [13]. We repeated our ultrastructural studies in heterothallic crosses of WT expressing Fus1-sfGFP and mCherry-D4H to unlabelled *gpd1Δ*, generating 32 tomograms for *h-gpd1Δ* x *h+* WT and 10 tomograms for *h*- WT x *h+ gpd1Δ* (Fig 7A-B, S7A). The frequency of membrane waviness was not substantially altered in *h+ gpd1Δ* pre-fusion cells (62.5%; 5/8) and slightly increased in *h*- cells (64.5%; 20/31) (Fig S7B), and the waves were more prominent than in WT cells. We confirmed by fluorescence microscopy that asymmetries in exo- and endocytosis were not altered in the *gpd1△* cells: indeed, the exocytic vesicle-associated Myo52 signal was higher and the endocytic vesicle-associated Fim1 signal lower in *h+* cells, as in WT cells (Fig 7C-D). These data are consistent with a reduction in turgor pressure in *gpd1△* cells exacerbating the membrane waviness phenotype.

**Figure 7.**
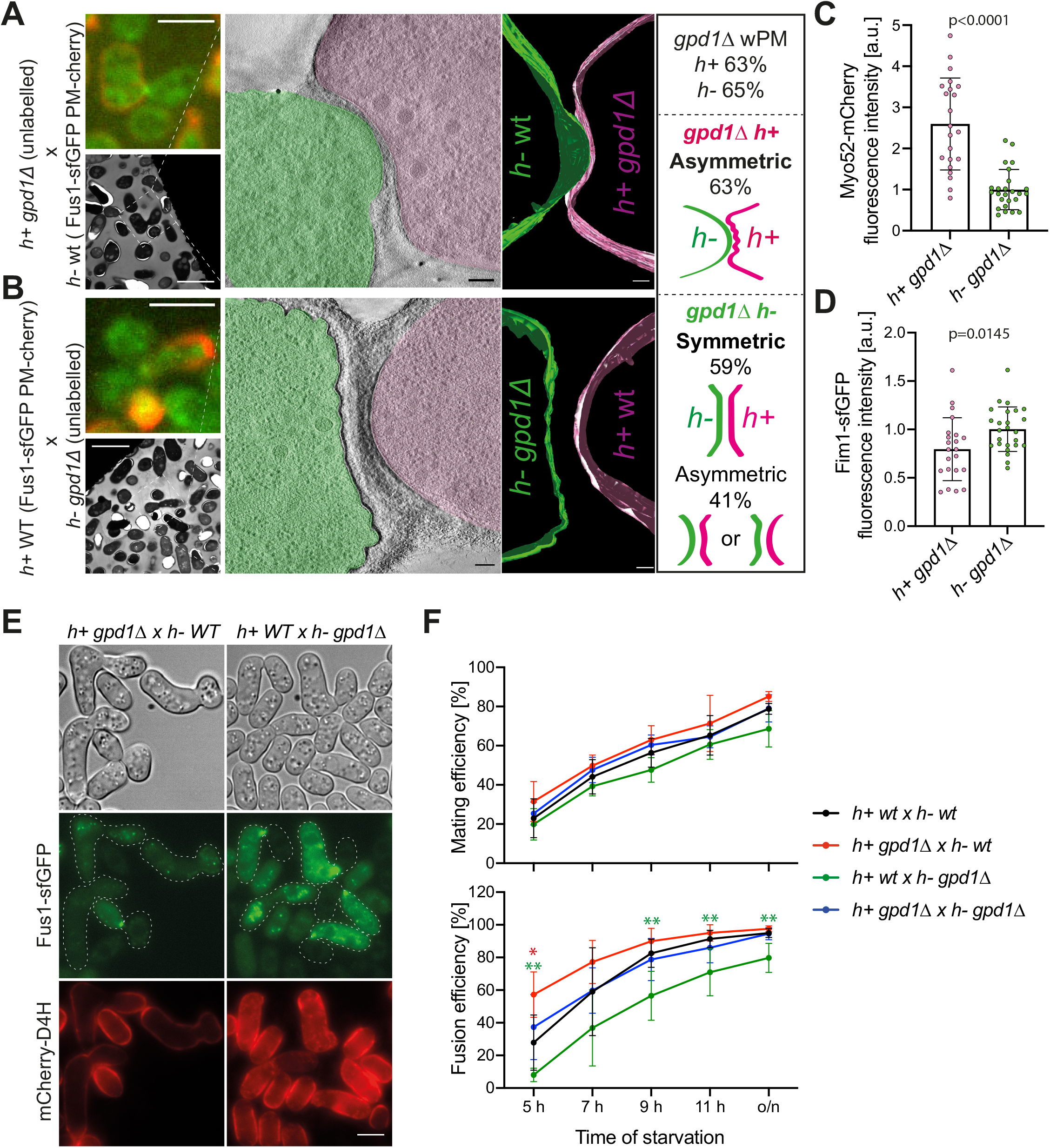
Different turgor pressure between cells of a pair is required for fusion. **(A-B)** Representative example of h-WT cells crossed with h+ *gpd1△* cells. The top-left light microscopy pictures show Fus1-sfGFP (green) and mCherry-D4H (red) expressed in the WT cell. Bottom-left low magnification TEM picture show correlation with the same mating pair. The virtual tomogram slices shows PM shapes, segmented with Imod. Scale bars are 5 μm in light microscopy, 10 μm in low magnification TEM pictures and 100 nm in tomogram slices. **(C-D)** Myo52-mCherry and Fim1-sfGFP mean fluorescence intensities of the 5 maximum pixels around each peak from profiles along the shmoo tips of tagged *gpd1△* cells crossed to WT imaged as in Fig 6E-F (n=22 h+ and 25 h-gpd1△ cells from 2 experiments; p-values are from Mann-Whitney tests). **(E)** Light microscopy images of WT x *gpd1△* crosses at 7h of nitrogen starvation. Green channel is Fus1-sfGFP and red channel is mCherry-D4H, both present only in WT cells. gpd1Δ cells are unlabelled. **(F)** Quantification of mating and fusion efficiencies over time (n ≥ 900 cells of each genotype from 5-7 experiments; * indicates p < 0.05; ** indicates p < 0.01 from unpaired t-tests).

Regarding membrane curvature, cell pairs in which the *h+* cell carried the *gpd1* deletion did not appear significantly different from WT. While our dataset is small, we recovered examples across the fusion process, which showed the same tendencies as WT pairs: both far-CW-contact cases were symmetric and of the 6 close-CW-contact cases (PM distance = 47.5 ± 30.7nm), 2 showed slight asymmetry and 3 had the *h*- cell protrude into a concave mutant *h+* (Fig S7C). Thus, further decrease of turgor pressure in *h+* does not alter, and may even accentuate, the general asymmetry between pre-fusion partner cells.

By contrast, when the *h*- cell carried the *gpd1* deletion, most cell pairs showed a symmetric (16/31) or only slightly asymmetric (8/31) curvature configuration. The remaining pairs showed a clearer asymmetry (7/31), but we did not find any example of a cell protruding into its partner. Furthermore, amongst all asymmetric situations, both mating types had the same probability of adopting the convex shape (7 *h+* vs 8 *h*-; Fig S7C, Fig 7B). We also noted that the vast majority of cell pairs showed substantial amount of cell wall between cells, with only 5 tomograms with minimal PM distance between cells <100 nm. However, the *h-gpd1Δ* x *h+* WT pairs did not look like the WT far-CW-contact class, as the cell walls of the two cells had merged together over a large surface area, leading to extended contact zones between cells. This suggests that the fusion process is delayed in these cell pairs. While fusion delay may reduce the chances of observing asymmetric configurations, we note that the pairs with cells closer than 100 nm, including one with PM contact, were also symmetric or only very slightly asymmetric. These data thus suggest that reduction in turgor pressure in *h*- cells prevents protrusion into the partner cell and leads to a more symmetric arrangement between partners.

To further investigate the importance of cell pair asymmetry on fusion success, we quantified mating and fusion efficiencies of *gpd1△* cells by light microscopy. Deletion of *gpd1* in one or both mating types did not affect the ability of cells to pair together. By contrast, deleting *gpd1* in one or the other cell type caused opposite effects on fusion efficiency (Fig 7E-F): when the *h*- cell lacked *gpd1*, which we showed above results in symmetric configurations, fusion was delayed; when the *gpd1* was deleted in the *h+* cell, which preserves or accentuates asymmetry, fusion occurred faster than in WT pairs. Importantly, when both partner cells lacked *gpd1*, fusion efficiency was restored to wildtype levels, indicating that the changes in fusion efficiency observed upon deleting *gpd1* in only one partner cell reflect the difference in turgor pressure between the two cells rather than an absolute requirement of *gpd1* in either cell type. We conclude that asymmetries in turgor pressure between cell types promote cell fusion.

## Discussion

By using correlative light-electron tomography, we reveal ultrastructural details of the fission yeast cell fusion process. While mating yeast cells are considered isogametes, which cannot be distinguished morphologically, our study now reveals that these cells display asymmetries at the ultrastructural level: one cell, generally the M-partner, protrudes into the other cell, which usually exhibits a less tensed plasma membrane. Tomograms of cell pairs at or around the time of membrane merging further suggest how fusion pores form and expand during cell fusion of a walled species.

### Organization of the fusion focus

The tomographic information reveals the general organization of the fusion focus as dense assembly of vesicles interspersed with linear filaments, largely excluding other organelles and ribosomes. These observations are consistent with the localization of the formin Fus1, actin-associated proteins, myosin V Myo52 and other secretory vesicle-associated markers by light microscopy [15, 17, 18]. The filaments very likely represent linear actin filaments, as the frequency of detection was reduced in cells lacking Fus1. We note that these mutant cells contain other actin nucleators, including the formin For3, so complete absence of filaments is not expected. There are two sub-populations of vesicles. The identity of the lower abundance, smaller-size vesicles is not certain: they may represent endocytic vesicles, but we cannot exclude that they represent a mixed population with other transport carriers. By contrast, we are highly confident that the most abundant population, with diameters averaging 60 to 70 nm, up to 100 nm, represents secretory vesicles. Indeed, this population is strongly diminished in cells lacking the formin Fus1, which concentrates the delivery of secretory markers at the cell growth projection [15]. Interestingly, the size of these vesicles is smaller than that reported during mitotic growth, where the average diameter of secretory vesicles is about 100 nm [26–28]. A similar observation was made in *S. cerevisiae*, where 60-70 nm vesicles accumulate at mating projections and fusion sites [22, 29], but 100 nm vesicles are reported in mutants of the secretory pathway during vegetative growth [30, 31]. Thus, secretory vesicles have a reduced size, and perhaps specific contents including cell wall digestive enzymes, during sexual reproduction.

### Formation and expansion of the fusion pore

Examination of the fusion pore revealed three notable features. First, the vast majority of fusion pores captured in our tomograms were large, generally extending beyond the tomographic volume. Indeed, only three pores were fully captured within the tomographic volume. This suggests that early stages of membrane fusion and pore expansion are rapid relative to other steps. However, we note that the existence of mutants that exhibit transient fusion indicates that this step is not absolutely unidirectional [32]. Second, we noticed that the edges of fusion pores often did not appear circular. In larger, more expanded pores, this manifested by the presence of a strand of cell wall penetrating into the pore. In tomograms that captured the edge of a pore along the plane of the section, these edges were often irregular with finger-like extensions into small regions lacking cell wall densities. We also identified such regions not associated with larger pores, suggesting these represent early stages of fusion, though the resolution limits imposed by the post cryo-fixation resin embedding do not allow to unambiguously trace both leaflets of the PM around these features. Third, we identified one tomogram, which displayed at least two such connections distant from each other, suggesting that membrane fusion can initiate at several positions. While merging of several expanding fusion pores may be the cause of the irregular edges, a second non-mutually exclusive possibility is that the rate of cell wall digestion limits the expansion of the fusion pore (Fig 3F). As cell wall digestion depends on secreted enzymes [15, 33–35], individual fusion events of secretory vesicles with the plasma membrane may lead to non-homogeneous enzyme distribution and cause degradation to occur at different rates around the pore edge.

### Control of plasma membrane tension during yeast mating

A striking feature of the plasma membrane in pre-fusion cells is its wavy pattern. This pattern is spatially restricted and strongly correlates with local exocytosis. Indeed, PM waves are specifically observed at the site of cell-cell contact, where membrane turnover through exo- and endocytosis is taking place, and are not seen in the lateral PM regions captured in our tomograms. The waves are shallower than Ω-shapes formed immediately upon vesicle-PM fusion, but the observation that they are almost completely abolished in *fus1△* cells argues that this pattern results from strong local secretion activity. In *S. cerevisiae*, undulating PM was also noted at the tips of mating projections [22] and can be seen in electron micrographs of fusion pairs (see for instance Fig 2H in [36]), suggesting that this is a conserved feature associated with strong local secretion during yeast sexual reproduction. The wavy pattern indicates reduced membrane tension, which does not instantaneously equilibrate with lateral membrane regions.

Interestingly, the occurrence of the wavy membrane pattern increased through the fusion process up to membrane merging and was more frequent in the P-cell partner. This correlates with previous measurements showing increase in Myo52 signal levels in the course of fusion [15] and with differences in the ratio of exo-to endocytic vesicles between cell types, where this ratio is higher in P than M-cells, as measured on both EM and light microscopy images. Exo- and endocytosis are generally strongly correlated and mutually influence each other through changes in membrane tension, which complicates the dissection of the cause of the cell type-specific difference. Indeed, studies in a large number of cell types have shown that high membrane tension stimulates exocytosis, which in turn leads to tension reduction [37]. Conversely, endocytosis is inhibited by membrane tension and contributes to its increase [38]. We envisage two possible scenarios. First, P-cells may have a higher rate of secretion. Consistent with this view, the number of exocytic vesicles and associated markers is higher in P-cells, and this higher signal depends on Fus1, suggesting that the cell type-associated difference is inherent to the fusion focus. Previous data had also shown a faster turnover of Myo52 by FRAP analysis in P-cells [15]. However, this scenario does not explain the reduction in endocytic vesicle-associated Fim1 signal in P-cells. A second scenario is that the cell type-specific difference resides in regulation of endocytosis, where P-cells may have a lower maximal endocytic capacity. This may cause a membrane ‘traffic jam’, where endocytosis cannot keep up with the exocytic capacity, leading to a local reduction of membrane tension, visible in the membrane waves, and consequent decrease in the rate of exocytosis, resulting in excessive accumulation of secretory vesicles. The levelling of both endo- and exocytic signals in *fus1△* cells may result from reduction in exocytosis, bringing compensatory endocytosis levels away from the maximal capacity reached in wildtype cells. While further work will be required to distinguish between these possibilities, either scenario leads to the same overall conclusion that the balance between exo- and endocytosis is shifted towards higher exocytosis in P-cells, which causes a local loss of membrane tension at the cell front.

### Control of cell protrusion by turgor pressure

Our ultrastructural analysis revealed a strong asymmetry in membrane curvature at the site of cell-cell contact, which increases as cells near each other during cell wall thinning, and culminates shortly before membrane fusion, with one cell, generally the M-cell, protruding into its partner. The asymmetry is relaxed post-fusion, as only few pairs with fusion pore showed asymmetry. Protruding M-cells also showed a membrane invagination on either side of the protrusion, which was unresolved in our tomograms. While we do not know at present the cause of this structural feature, its frequency suggests it may be an important structure to help the organization of the fusion site.

We explored the hypothesis that turgor pressure, more specifically a difference in turgor between partner cells, may be at the origin of the protrusion. Indeed, deletion of Gpd1 in M-cells largely prevents their protrusion into P-cells and yields more symmetrical fusion pairs. By contrast, its deletion in P-cells does not prevent M-cell protrusion. Gpd1 functions in the glycerol biosynthetic pathway and, upon hyper-osmotic stress, serves to restore osmotic pressure through glycerol synthesis [24, 25]. During vegetative growth, deletion of Gpd1 reduces the growth force of cells placed in hyper-osmotic conditions [13]. During sexual reproduction, our results indicate that *gpd1△* cells have reduced growth force even without external hyper-osmotic challenge. This is consistent with Gpd1 expression, which is induced upon osmotic shock [39–41], but also increases about 6-fold during mating [11]. We note that strong turgor is not the only requirement for protrusion, as disruption of the fusion focus in *fus1Δ* likely also interferes with formation of the protrusion in M-cells though our dataset is small. However, our results strongly support the view that regulation of turgor pressure is required for the asymmetric protrusion of M into P-cells.

Regulation of turgor pressure is likely to be generally important to control the fusion of walled cells. In *S. cerevisiae*, cross-talks between HOG-MAPK signalling, which controls glycerol production in response to osmotic stress, and pheromone-MAPK signalling have been described, where each pathway downregulates the other [42–45]. The pheromone-MAPK pathway is also thought to indirectly promotes HOG-MAPK signalling, by stimulating glycerol release through the glycerol efflux aquaglyceroporin Fps1 and consequent HOG-dependent glycerol synthesis to compensate for turgor loss [46]. Thus, mating *S. cerevisiae* cells may have fast turgor adaptation through increased glycerol turnover. Interestingly, interfering with glycerol turnover through deletion of Fps1 in one, but not both, partner cell causes an increase in fusion intermediates, indicating a block or delay in fusion, though deletion of Gpd1 had no effect in this organism [47]. Whether this system operates differently in the two mating types has not been investigated, but interestingly, the deleterious effect of *fps1Δ* was stronger when the a-cell was mutant [47], suggesting a possible asymmetry between cell types.

### Origin of yeast gamete asymmetry

An open question is how the asymmetry between cells comes about. Our analysis clearly shows that ultrastructural asymmetry, for both membrane tension and curvature, is strongly linked with cell type: P-cells have a higher exo-to endocytic ratio leading to low membrane tension, and M-cells exhibit increased turgor pressure, which drives protrusion and likely also contributes to membrane tension increase. These cell type-linked differences are also visible at the functional level, as *gpd1Δ* is detrimental only in M-cells while the phenotype of *fus1Δ* is more severe in P-cells. However, our analysis of heterothallic wildtype cells also revealed three cases of asymmetry reversal, where the P-cell was protruding into the M-partner. This suggests that asymmetry is not completely hardwired and may be amplified from small initial differences between cell types.

At the transcriptional level, there are very few differences between mating types. Only 16 genes have been reported to be differentially expressed, most of which code for agglutinins, pheromones, processing enzymes and receptors [11, 12]. The origin of cell type-specific differences may reside in the different pheromone receptors: though acting through the same signalling pathway, they may elicit quantitatively different signals that alter crosstalk with the osmo-sensing pathway. Alternatively, turgor pressure differences may be linked to one uncharacterized small gene (SPAC1565.03) reported to be expressed specifically in P-cells [11, 12], and which was independently identified to promote growth on glycerol, like *gpd1* [48]. We note that *gpd1* expression itself, though induced during mating, is unlikely to be the key cell type-specific difference, as deletion of *gpd1* in both partner cells restores wildtype-like fusion kinetics and thus likely functional asymmetries. Because asymmetries increase during the course of fusion, this also suggests that they are amplified by cell-cell interaction either through increase in signalling or through mechanical signals upon contact with the partner cell.

### Function of membrane asymmetries

It is remarkable that cell fusion is more strongly compromised by *gpd1* deletion in the M-cell and *fus1* deletion in the P-cell. These situations reduce the asymmetries observed in wildtype cells, by preventing protrusion and membrane waviness, respectively. Because membrane waves are observed primarily in P-cells, we hypothesize that their loss is the cause of the stronger *fus1Δ* fusion defect in P-cells. However, as local secretion organized by the fusion focus is also necessary for cell wall digestion [15], we cannot exclude that the stronger effect on cell fusion upon deletion of *fus1* in the P-cell may result from a more important contribution of P-cells to cell wall digestion. Similarly, because it is primarily the M-cell that forms a protruding, convex shape, loss of this shape is the likely cause of the fusion delay when *gpd1* is deleted only in this cell. By contrast *gpd1* deletion in the P-cell, which maintains or exacerbates the protrusion asymmetry, leads to faster cell fusion. These observations strongly support the view not only that the two cells assume different roles, but also that the resulting plasma membrane asymmetries are important for efficient cell-cell fusion.

The asymmetry of yeast cell fusion recalls similar asymmetry observed in the fusion of *Drosophila* myoblasts, where fusion-competent myoblasts form a podosome that protrudes into the myotube [5]. Podosome-like structures are also observed in other types of somatic cell fusion [4]. Although the underlying protrusive force generation may be different [16], the similar configurations suggest that evolution of an asymmetric setup is beneficial to the success of cell-cell fusion. We hypothesize that membrane organization asymmetry directly promotes plasma membrane merging. Local membrane curvature caused by high local secretion in the P-cell may help destabilize the bilayer and lower the energy required to initiate membrane merging upon contact with the partner cell. The turgor pressure-dependent force causing the protrusion, which is initially countered by the intervening cell wall, may upon complete cell wall removal, help overcome repulsive forces to bring the two cells’ membranes close enough for merging. While the identity of a possible fusogenic machinery is still unknown, these ultrastructural plasma membrane features suggest that biophysical changes in membrane tension and curvature may be important contributors to the membrane fusion process.

## Materials and methods

### Strains and growth conditions

*S. pombe* strains used in this study are listed in Table S1. Heterothallic *h+* and *h*- strains and homothallic (*h90*) strains, able to switch mating types, were grown in Minimum Sporulation Media (MSL) supplemented with nitrogen (+N). Mating assays were conducted on MSL without nitrogen, essentially as described [49], except for *gpd1Δ* strains, which were mated on Malt Extract (ME).

### Strain construction

Strains were constructed using standard genetic manipulation of *S. pombe* either by tetrad dissection or transformation.

For generation of *fus1* deletion mutants, WT strains were transformed with a linearized deletion plasmid (based on the pFA6a-kanMX backbone) containing at least 400 bp of homology to *fus1* gene flanking regions (pSM1966). For generation of *gpd1* deletion mutants, the *gpd1* ORF was replaced by *ura4* ORF. This was achieved by PCR amplification of a fragment from strain NM204 (received from Fred Chang, UCSF) carrying *ura4+* ORF at the *gpd1* locus, which was transformed and integrated in the genome of *ura4-D18* strains by homologous recombination. For C-terminal tagging of *myo52* with mCherry, strains were transformed with linearized fluorophore tagging plasmid containing at least 400 bp of homology to the C-terminal part of the gene, in frame mCherry, kanMX resistance and *myo52* 3’UTR (pSM2735). For tagging fim1 with sfGFP, a pFA6a-sfGFP-natMX plasmid (pSM1686) was used as a template for PCR-based targeted tagging, as described [50].

### Mating Assays

For both light microscopy and CLEM experiments cells were first pre-cultured overnight in MSL+N at 25°C, then diluted to OD600 = 0.025 into MSL+N at 30°C for 16 hours. The amount of exponentially growing cells equivalent to OD600 of 3-5 (depending on the experiments but constant within the same experiment) were pelleted, washed in MSL-N by 3 rounds of centrifugation, and resuspended in 50-100 μl MSL-N. Cells were then placed in 10 μl drops on 2% agar MSL-N or 2% agar ME plates at 30°C to allow mating. Samples for CLEM were further processed after 5h (see CLEM and tomography section below). Samples for time-lapse live imaging were transferred to MSL-N 2% agar pads after 4h at 30°C and mounted on a microscope slide, covered with coverslip, sealed with VALAP, incubated one more hour on the pad at 30°C and imaged. Samples for experiments to measure mating and fusion efficiencies were placed on ME and maintained at 30°C for 5, 7, 9, 11 and 24 hours and cells were imaged at the indicated times after mounting them on slides and covered with coverslips.

### Light microscopy

Light microscopy images in figure 7C and S7D were obtained using wide-field microscopy performed on a DeltaVision platform (Cytiva, Massachusetts, USA) composed of a customized inverted microscope (IX-71; Olympus, Japan), a 100 × /1.4 NA oil objective, a camera (CoolSNAP HQ2; Photometrics, Arizona, USA, or 4.2Mpx PrimeBSI sCMOS camera; Photometrics, Arizona, USA), and a color combined unit illuminator (Insight SSI 7; Social Science Insights). Images were acquired using softWoRx v4.1.2 software (Cytiva, Massachusetts, USA). Images shown are single-plane views.

Images presented in figures 5C and 5D or used to get data for figures 6E, 6F, 7E, 7F and S5D are average projections of time-lapse movies acquired a maximum speed during 3 minutes using a spinning-disk microscope composed of an inverted microscope (DMI4000B; Leica) equipped with an HCX Plan Apochromat 100 × /1.46 NA oil objective (PerkinElmer; including a real-time confocal scanning head [CSU22; Yokagawa Electric Corporation], solid-state laser lines, and an electron-multiplying charge coupled device camera [C9100; Hamamatsu Photonics]). Images were acquired using the Volocity software (PerkinElmer).

For CLEM, light microscopy images for the WT h90 samples were acquired on a Nikon TE2000-E microscope equipped with a 100x 1.49 NA TIRF oil objective, a NEO sCMOS DC-152Q-C00-FI camera (Andor) and a Lambda DG-4 lamp (Sutter Instruments). Images were acquired using the NIS Elements software (Nikon). All other images were acquired on the DeltaVision platform described above, but with a UPlan Apochromat 60 × 1.42 NA oil objective.

### CLEM and tomography

CLEM was as described in [51], adapted for mating cells. Briefly, cells were grown for mating as described above. After washes to remove nitrogen cells were added into MSL-N plates and ME plates for *gpd1* deletion mutant strains. We allowed cells to mate for 5 h. A few μl of MSL-N was pipetted onto the cells to form a thick slurry, which was pipetted onto a 3-mm-wide, 0.1-mm-deep specimen carrier (Wohlwend type A) closed with a flat lid (Wohlwend type B) for high-pressure freezing with a HPM100 (Leica Microsystems) (for *h90* samples) or a Wohlwend HPF Compact 02. The carrier sandwich was disassembled in liquid nitrogen before freeze substitution. High-pressure frozen samples were processed by freeze substitution and embedding in Lowicryl HM20 using the Leica AFS 2 robot as described [51]. 300 nm sections were cut with a diamond knife using a Leica Ultracut E or Ultracut UC7 ultramicrotome, collected in H_2_O, and picked up on carbon-coated 200 mesh copper grids (AGS160; Agar Scientific). For light microscopy, the grid was inverted onto a 1× PBS drop on a microscope coverslip, which was mounted onto a microscope slide and imaged as indicated above. The grid was then recovered, rinsed in H_2_O, and dried before post-staining with Reynolds lead citrate for 10 min. 15-nm protein A–coupled gold beads were adsorbed to the top of the section as fiducials for tomography. Transmission electron micrographs (TEM) were acquired on a FEI Tecnai 12 at 120 kV using a bottom mount FEI Eagle camera (4kx4k). Low magnification images were acquired at 17.816 nm pixel size and high magnification at 1.205 nm pixel size. A few TEM acquisitions (including panel 3D) were performed on a TF20 microscope (FEI) in STEM mode, with an axial brightfield detector, using a camera length of 200 mm and a 50 μm C2 aperture [52]. For tomographic reconstruction of regions of interest, tilt series were acquired at 1.205 nm pixel size over a tilt range as large as possible up to +/-60° at 1° increments using the Serial EM software [53]. IMOD software package with gold fiducial alignment [54, 55] was used for tomogram reconstruction, segmentation and modelling.

### Quantification and Statistical Analysis

Minimal distance between PMs, and density and diameters of vesicles in tomograms where manually measured within a cylinder of diameter 1 μm centred at the centre of the cells’ contact zone using IMOD software. Vesicle measurements were done for each cell separately in the half-cylinder covering that cell.

Myo52-tdTomato or mCherry, Fim1-sfGFP and Exo84-GFP fluorescence profiles at shmoo tips were obtained on average projections of time-lapse movies acquired at maximum speed during 3 minutes. Fluorescence intensity along a 7-pixel-wide segmented line was collected using the FIJI plot profile tool. Values were corrected for the external background. The curves were centered on the maximum pixel values of the Myo52 channel. Values in figures 5F, 6E (right), 6F (right), 7E, 7F and S5F were obtained from the average of the 5 pixels around each peak maximum from the profiles described above.

Mating and fusion efficiencies were calculated as in [15]: mating efficiency represents the fraction of cells engaged in mating; fusion efficiency represents the fraction of mating pairs that have fused. Mating pairs were identified by the presence of Fus1-sfGFP or Myo52-tdTomato at the fusion focus. The transfer of mCherry-D4H or Fimbrin1-sfGFP into the unlabelled partner was used to identify fused pairs.

Statistical analysis was performed using GraphPad Prism. Statistical significance was determined using one-way ANOVA test, paired or unpaired t-test or Mann-Whitney test, as indicated.

## Acknowledgements

We thank the University of Lausanne EM facility (EMF), in particular Jean Daraspe for help with electron microscopy sample preparation and acquisition, Bruno Humbel for help with manipulation of the electron microscope and Christel Genoud for discussions and suggestions. We thank Fred Chang for sharing a *gpd1Δ* strain and xxx for comments on the manuscript. This work was funded by an ERC Consolidator grant (CellFusion) and a Swiss National Science foundation grant (310030B_176396) to SGM, and a “poste de soutien” awarded to OML. WK was supported by the Medical Research Council (MC_UP_1201/8).

## Author contribution

SGM prepared CLEM samples and reconstructed tomograms for the *h90* cells with help from WK. OML acquired EM data for all samples except for a few acquired by WK, and prepared CLEM samples and reconstructed tomograms for all heterothallic crosses. OML performed all other experiments with technical support from LM. OML analyzed the data together with SGM. OML and SGM prepared figures and wrote the manuscript, with comments from WK. OML and SGM acquired funding.

---

**Supplementary Figure 1.**
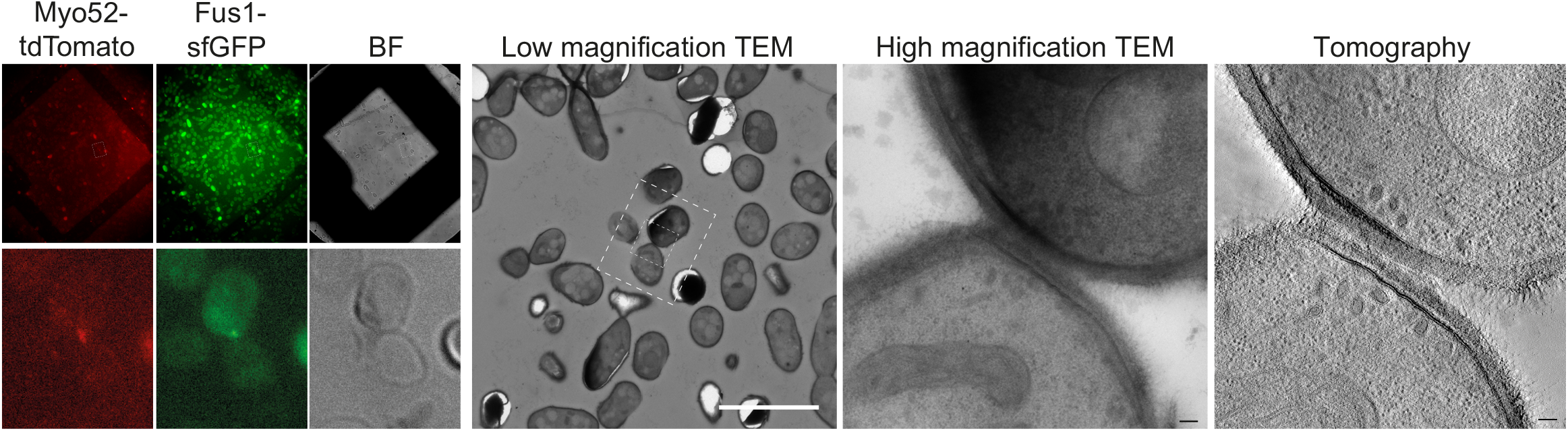
Example of correlation between light and electron microscopy during mating. Light microscopy images are GFP and tdTomato signals on 300nm sections of resin-embedded yeast cells expressing Fus1-GFP and Myo52-tdTomato, placed on EM grids. Size of grid square is 90.5 μm. Cells with fusion focus fluorescence signal are identified in low magnification (17.816 nm pixel size) TEM according to their position, preservation is tested at high magnification (1.205 nm pixel size) TEM, and tilt series are acquired at high magnification (1.205 nm pixel size) and processed for tomographic 3D reconstruction. Scale bars are 10 μm in low magnification TEM and 100 nm in high magnification TEM and virtual z-slice through tomogram.

**Supplementary Figure 2.**
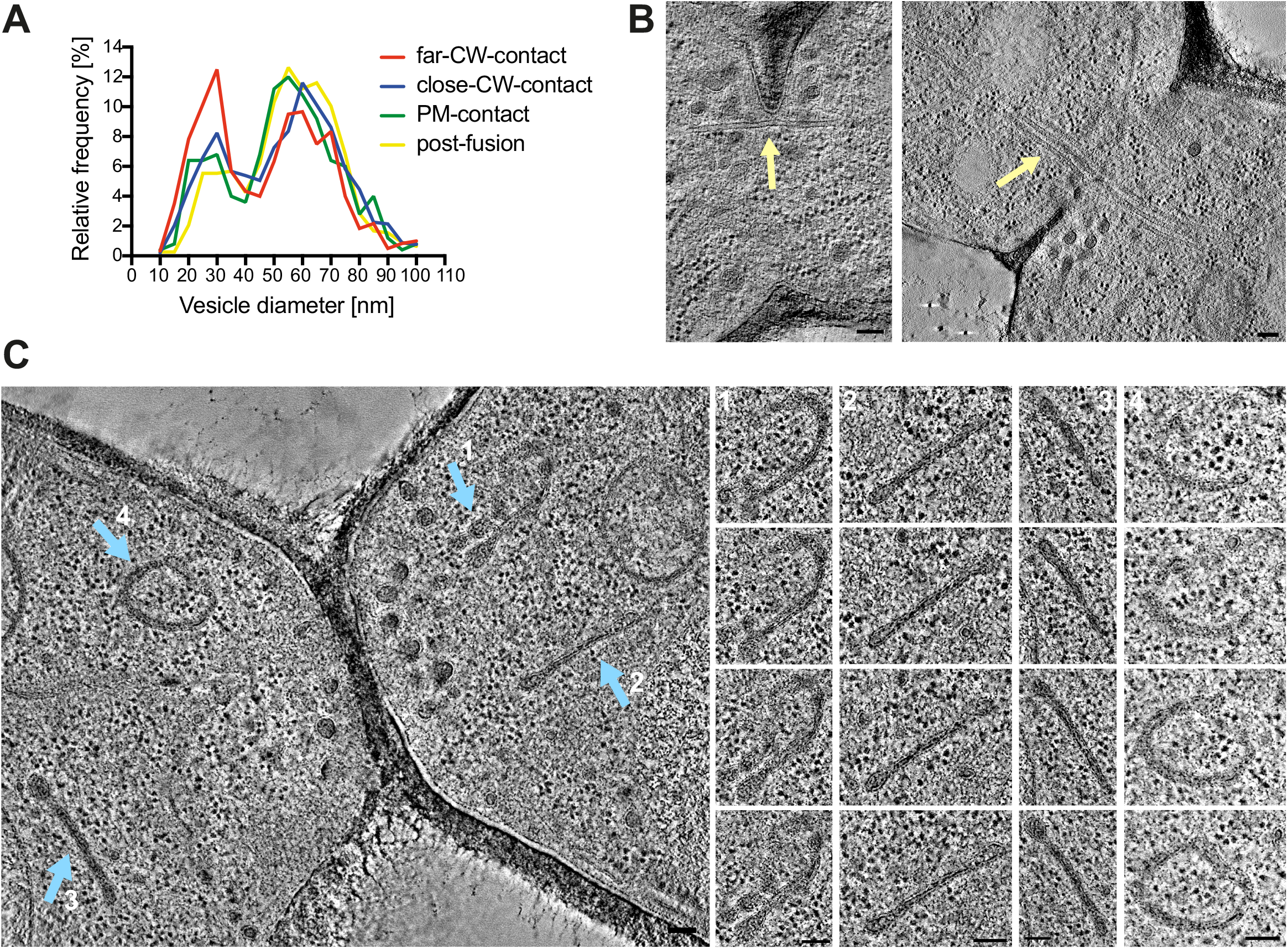
Ultrastructural features at sites of yeast cell fusion. **(A)** Size distribution of vesicles separated in the four stages: far-CW-contact (n=20 cells; 616 vesicles), CW-contact close (n=60 cells; 1865 vesicles), PM-contact (n=10 cells; 252 vesicles) and post-fusion (n=16; 788 vesicles). Bin width = 5 nm. **(B-C)** Virtual z-slices through electron tomograms of mating cells showing microtubule bundles crossing the fusion pore (B, yellow arrow) and organelles with similar density to secretory vesicles, organized in sheets or reticulated structures (C, blue arrows). Scale bars 100 nm.

**Supplementary Figure 3.**
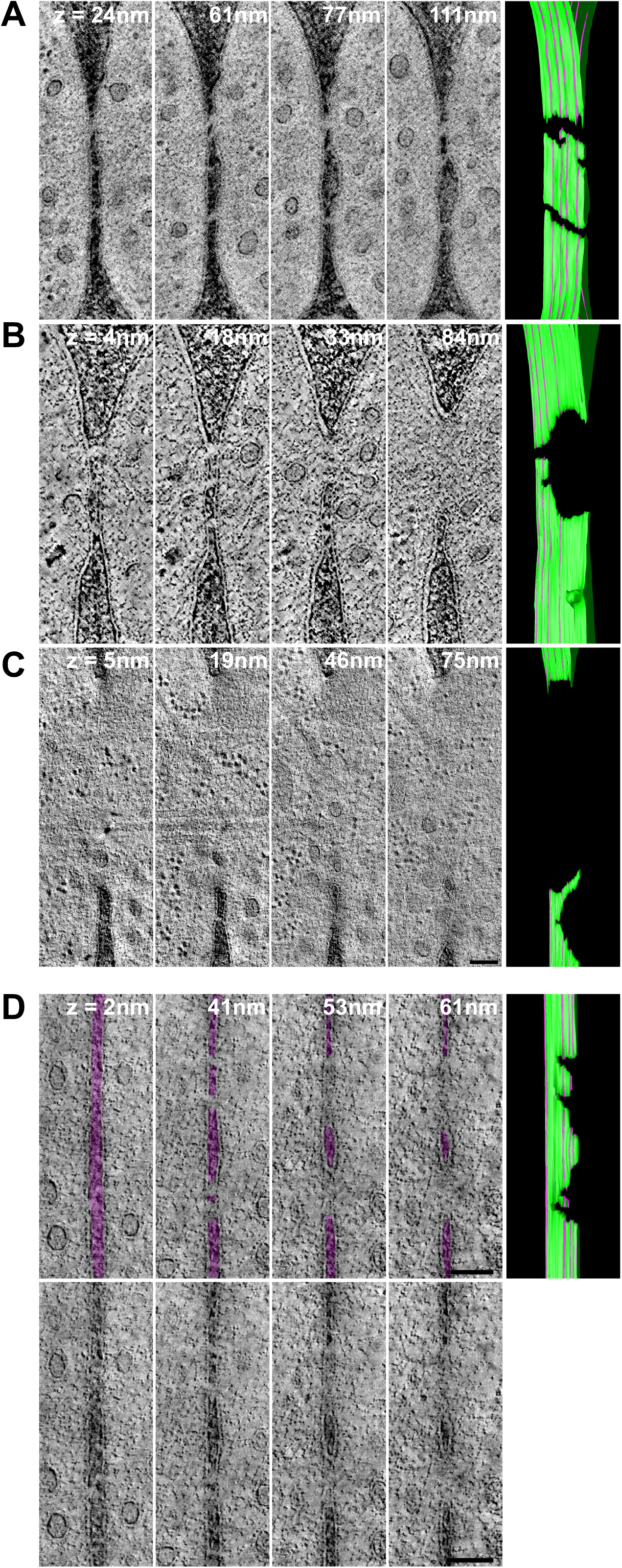
Shape of the fusion pore. **(A-C)** Original non-coloured virtual z-slices through electron tomograms showed in Fig 3C-D. **(D)** Four virtual z-slices through an electron tomogram showing two large pores with irregular edges. These two pores are likely joined in a single pore outside the reconstructed tomographic volume. On the top line, the cell wall is coloured in transparent purple to help visualization. Corresponding models of the outer leaflet of the PM are on the right. The location of each z-slice is shown by the purple lines on the model. Scale bars 100 nm.

**Supplementary Figure 4.**
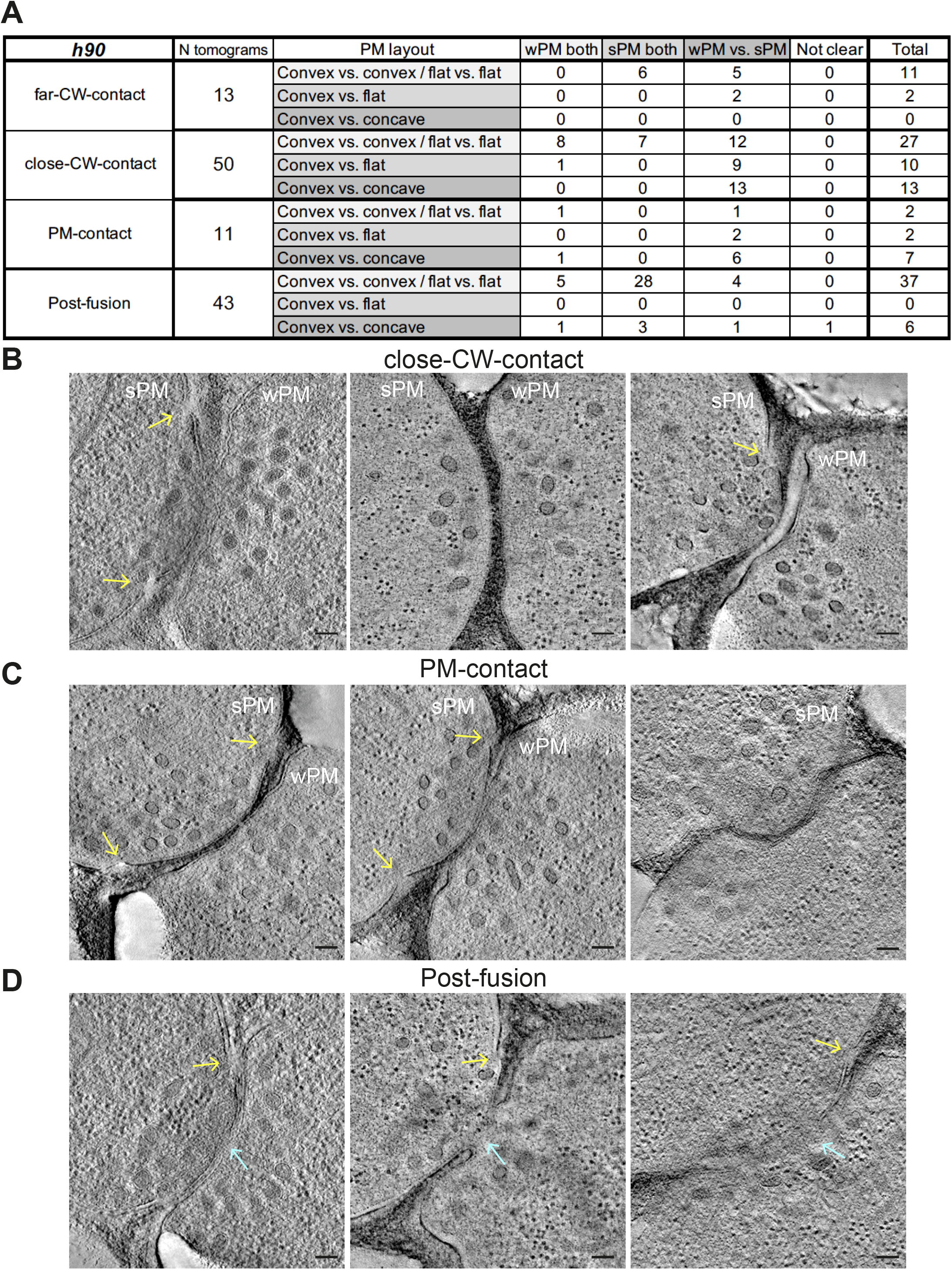
PM asymmetries during cell fusion. **(A)** Table classifying the set of tomograms from h90 WT cells used in this study, according to PM morphology along the four stages of fusion. **(B-D)** Virtual z-slices through electron tomograms of mating cells showing PM asymmetries (convex versus concave) in close-CW-contact stage (B), PM-contact stage (C) and post-fusion stage (D). Blue arrows point to fusion pores; yellow arrows point to unresolved regions at the PM). Scale bars 100 nm.

**Supplementary Figure 5.**
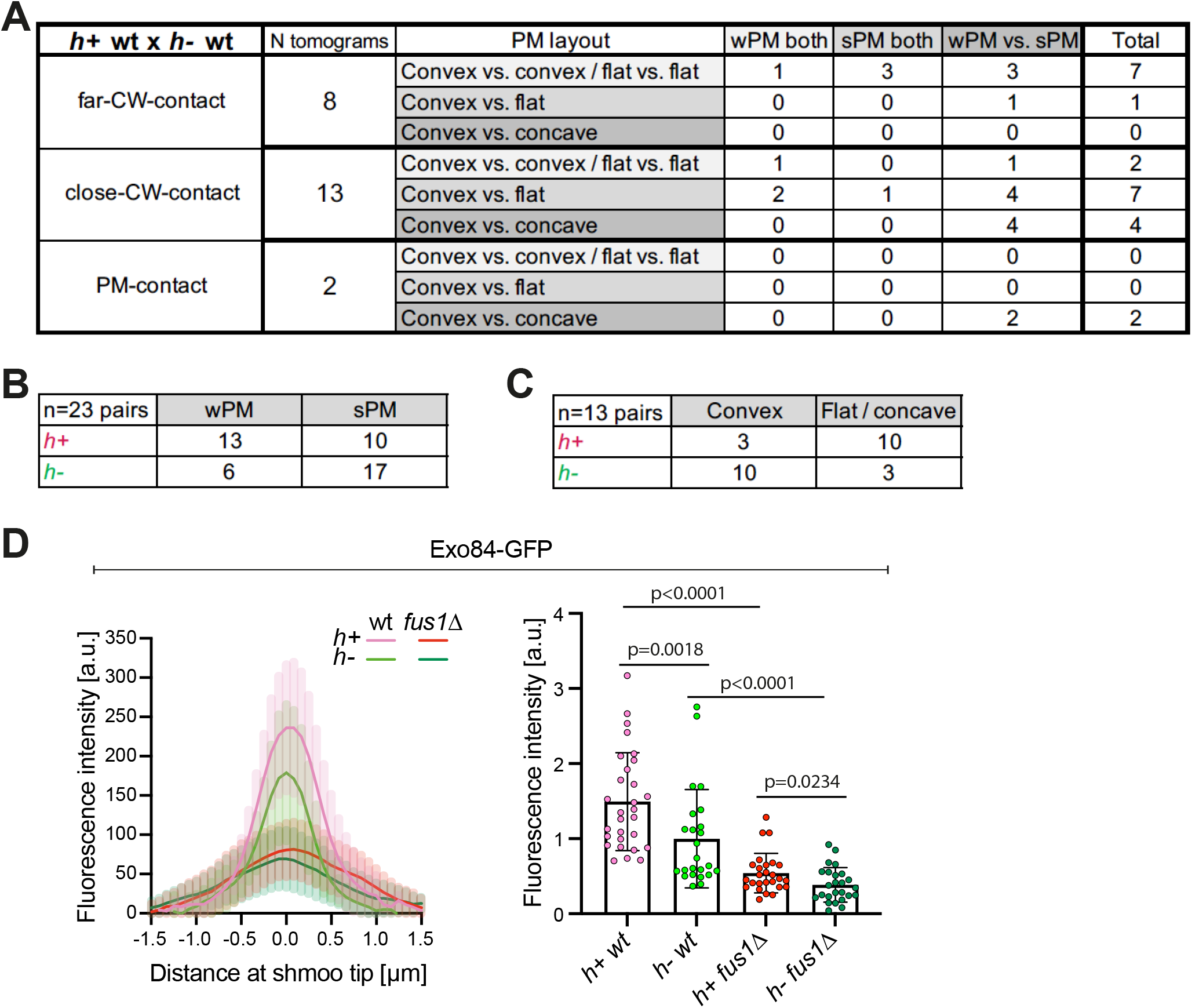
Mating type-specific plasma membrane asymmetries and concentration of exocytic vesicles. **(A)** Table classifying the set of tomograms from WT h+ x h-cell pairs used in this study, according to PM morphology. **(B)** Table showing the number of each mating type with wPM or sPM. **(C)** Table showing, amongst asymmetric mating pairs, the number of each mating type with convex or flat/concave shmoo tip conformation. **(D)** (Left) Fluorescence profile of Exo84 (GFP) along the shmoo tips of labelled WT and *fus1△* cells crossed to unlabelled WT cells. Images were obtained through average projections of time-lapse movies acquired a maximum speed during 3 minutes (n=16 h+ WT cells, 13 h-WT cells, 15 h+ *fus1△* cells and 14 h-*fus1△* cells). (Right) Exo84 mean fluorescence intensity of the 5 maximum pixels around peak from profiles as the left (n=29 h+ WT cells, 24 h-WT cells, 26 h+ *fus1△* cells and 25 h-*fus1△* cells from 2 experiments; p values from Mann-Whitney tests).

**Supplementary Figure 6.**
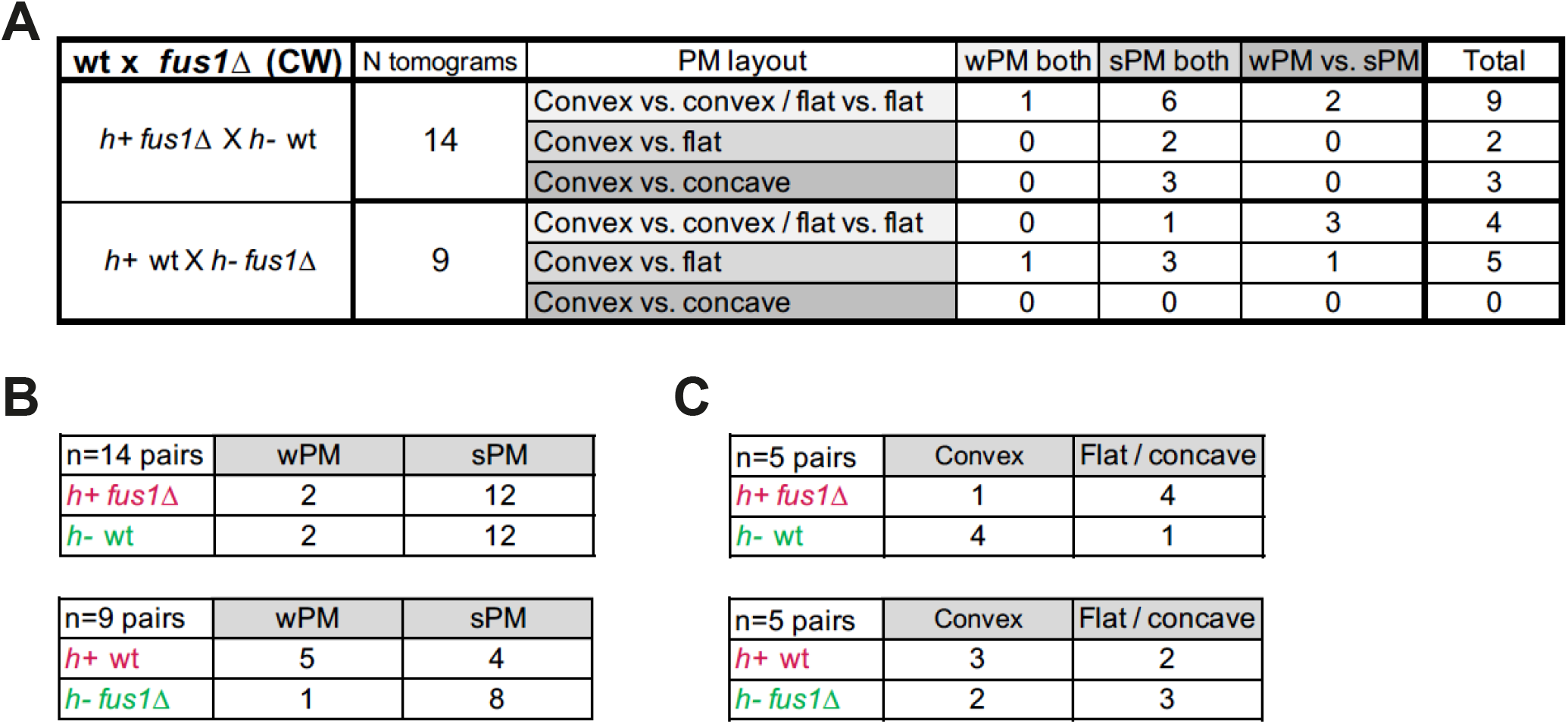
Classification of WT x *fus1*△ tomograms. **(A)** Table classifying the set of tomograms from WT x *fus1Δ* crosses used in this study, according to PM morphology in the CW-contact stage. **(B)** Tables showing the number of each mating type with wPM or sPM. **(C)** Tables showing, amongst asymmetric mating pairs, the number of each mating type with convex or flat/concave shmoo tip conformation.

**Supplementary Figure 7.**
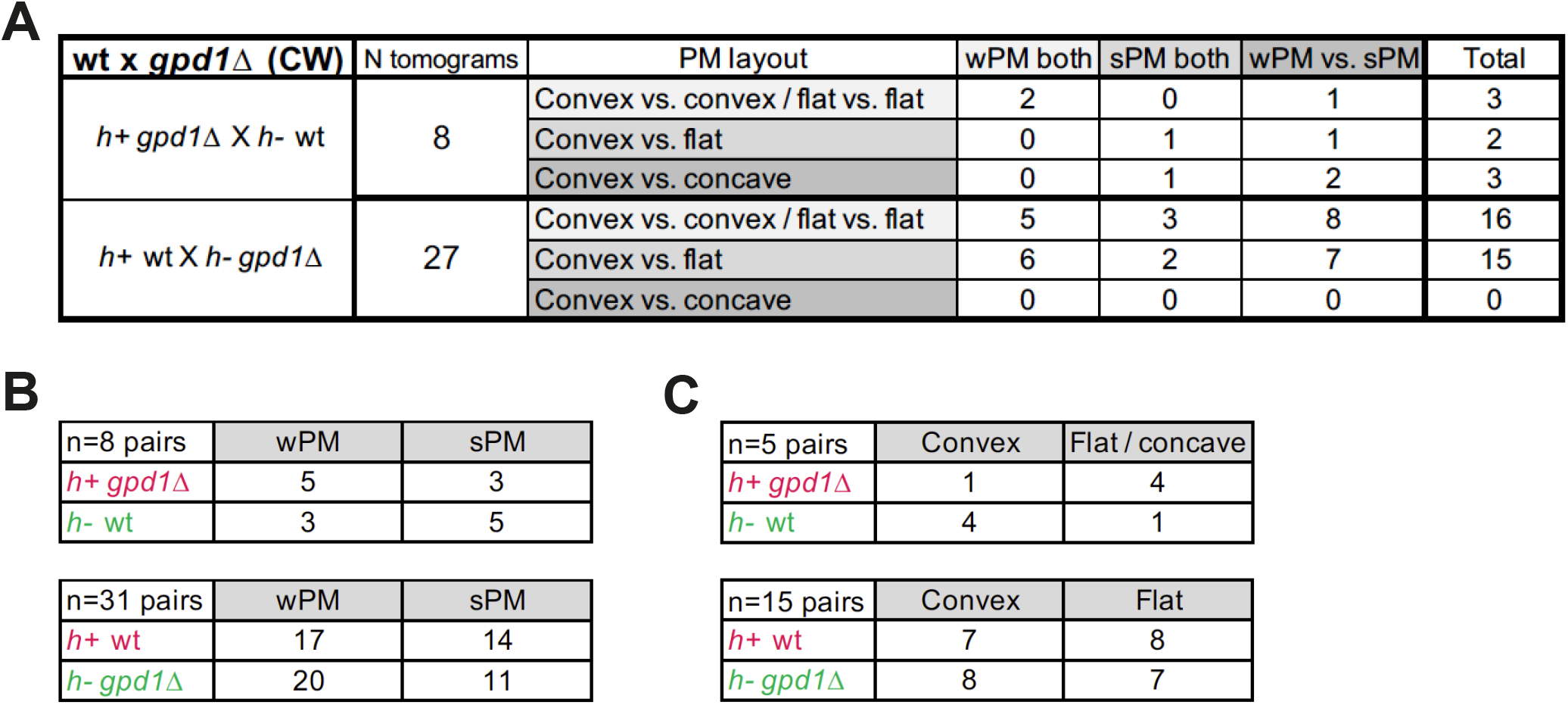
Classification of WT x *gpd1*△ tomograms. **(A)** Table classifying the set of tomograms from WT x *gpd1△* crosses used in this study, according to PM morphology in the CW-contact stage. **(B)** Tables showing the number of each mating type with wPM or sPM. **(C)** Tables showing, amongst asymmetric mating pairs, the number of each mating type with convex or flat/concave shmoo tip conformation.

**Supplementary Table 1:** Yeast strain used in this study.

| Strain | Source | Identifier |
| --- | --- | --- |
| *h+ WT* | Lab stock | YSM1371 |
| *h- WT* | Lab stock | YSM3904 |
| *h- WT* | Lab stock | YSM1372 |
| *h- exo84-GFP:kanMX myo52-tdTomato:natMX* | Lab stock | YSM3886 |
| *h+ exo84-GFP:kanMX myo52-tdTomato:natMX* | Lab stock | YSM3887 |
| *h90 myo52-tdTomato:natMX fus1-sfGFP:kanMX* | This work | YSM3888 |
| *h+ fus1-sfGFP:kanMX D4H-mCherry:ura4+ ura4-294* | This work | YSM3889 |
| *h- fus1-sfGFP:kanMX D4H-mCherry:ura4+ ura4-294* | This work | YSM3890 |
| *h- fim1-sfGFP:kanMX myo52-tdTomato:natMX* | This work | YSM3891 |
| *h+ fim1-sfGF: kanMX myo52-tdTomato:natMX* | This work | YSM3892 |
| *h+ fus1Δ::kanMX* | This work | YSM3893 |
| *h- fus1Δ::kanMX* | This work | YSM3894 |
| *h+ fus1Δ::hphMX fim1-sfGFP:kanMX myo52-tdTomato:natMX* | This work | YSM3895 |
| *h- fus1Δ::hphMX fim1-sfGFP:kanMX myo52-tdTomato:natMX* | This work | YSM3896 |
| *h+ gpd1Δ ::ura4+ ura4-D18* | This work | YSM3897 |
| *h- gpd1Δ::ura4+ ura4-D18* | This work | YSM3898 |
| *h+ fim1-sfGFP:kanMX myo52-tdTomato:natMX mTagBFP2:ura4+* | This work | YSM3899 |
| *h+ gpd1Δ::ura4+ fim1-sfGFP:NatMX myo52-mCherry:KanMX ura4-D18* | This work | YSM3900 |
| *h- gpd1Δ::ura4+ fim1-sfGFP:NatMX myo52-mCherry:KanMX ura4-D18* | This work | YSM3901 |
| *h+ fus1Δ::hphMX exo84-GFP:kanMX myo52-tdTomato:natMX* | This work | YSM3902 |
| *h- fus1Δ::hphMXexo84-GFP:kanMX myo52-tdTomato:natMX* | This work | YSM3903 |

## Movie legends

**Movie 1. Tomogram and model of a cell pair pre-fusion.**

The movie shows all virtual z-sections of the tomographic reconstruction of a pre-fusion site at the close-CW-contact stage. The plasma membrane is segmented in cyan, vesicles are shown in pink (large vesicles) and purple (small vesicles), and filaments are segmented with a yellow line. The left cell shows a convex shape with a smooth plasma membrane. The cell on the right shows a slightly concave shape at the site of cell-cell contact with plasma membrane that exhibits an undulating pattern.

**Movie 2. Tomogram and model of a cell pair post-fusion.**

The movie shows all virtual z-sections of the tomographic reconstruction of a cell pair postfusion. The plasma membrane is segmented in cyan, vesicles are shown in pink (large vesicles) and purple (small vesicles), and filaments are segmented with a yellow line.

**Movie 3. Tomogram and model of a cell pair with several connections.**

The movie shows all virtual z-sections and plasma membrane segmentation of the tomographic reconstruction of a cell pair exhibiting at least two apparent connections between cells. This corresponds to the example shown in Fig 3C and S3A.

**Movie 4. Tomogram and model of a cell pair with an asymmetric pore.**

The movie shows all virtual z-sections and plasma membrane segmentation of the tomographic reconstruction of a cell pair with a non-circular pore. This corresponds to the example shown in Fig 3D and S3B.

**Movie 5. Tomogram and model of a cell pair with an asymmetric pore.**

The movie shows all virtual z-sections and plasma membrane segmentation of the tomographic reconstruction of a cell pair with a non-circular pore. This corresponds to the example shown in Fig S3D.

**Movie 6. Tomogram and model of a cell pair with a pore containing a strand of cell wall.**

The movie shows all virtual z-sections and plasma membrane segmentation of the tomographic reconstruction of a cell pair whose fusion pore is split in two by a strand of cell wall. This corresponds to the example shown in Fig 3E and S3C.

